# Cooperative Control of Arrestin Activation By Membrane Lipids And Phosphorylation Barcodes

**DOI:** 10.64898/2026.04.13.718317

**Authors:** Yasmin Aydin, Ya Zhuo, Yu-Chen Yen, Chun-Liang Chen, Candice S. Klug, Adriano Marchese, Qiuyan Chen

**Affiliations:** Department of Biochemistry, Molecular Biology and Pharmacology, Indiana University School of Medicine, Indianapolis, IN 46202, USA; Department of Biochemistry, Medical College of Wisconsin, Milwaukee, WI 53226, USA; Department of Molecular Biosciences, University of South Florida, Tampa, FL 33620, USA; Moffitt Cancer Center, Tampa, FL 33612, USA; Department of Biophysics, Medical College of Wisconsin, Milwaukee, WI 53226, USA; Indiana University Melvin and Bren Simon Comprehensive Cancer Center, Indianapolis, IN 46202, USA

## Abstract

Arrestins regulate G protein–coupled receptor (GPCR) signaling by binding phosphorylated receptors embedded in lipid bilayers, yet how receptor phosphorylation and membrane composition cooperate to control arrestin activation remains unclear. Here, we reconstitute this interplay using N-terminally palmitoylated phosphopeptides tethered to nanodiscs of defined lipid composition and quantitatively measure arrestin-2 (Arr2) activation and membrane engagement. We find that both receptor phosphorylation and the lipid environment are essential for robust Arr2 activation, with phosphoinositides (PIPs) and other anionic lipids facilitating Arr2 activation and membrane association through distinct mechanisms. Systematic profiling of phosphorylation barcodes derived from atypical chemokine receptor 3 (ACKR3) and vasopressin receptor 2 (V_2_R) identifies phospho-motifs that potently activate Arr2. Moreover, the position of these motifs relative to the membrane determines Arr2 engagement, supporting a model of regional phosphorylation barcodes. Genome-wide motif analysis further links the phosphorylation barcode to predicted arrestin coupling strength and classification into Class A or Class B GPCRs. Finally, lipidated phosphopeptides inhibit GPCR–Arr2 interactions in live cells and enable structural characterization of Arr2–phosphopeptide complexes by cryo-electron microscopy, establishing a membrane-integrated framework for decoding arrestin response.

## Introduction

The non-visual arrestins, arrestin-2 (Arr2) and arrestin-3 (Arr3), are versatile adaptor proteins that mediate desensitization, internalization, and signaling across the G protein–coupled receptor (GPCR) superfamily^1,2^. Despite the remarkable diversity of GPCRs, these two arrestins effectively regulate downstream signaling by integrating receptor-derived cues, including phosphorylation patterns, receptor activation states, and the surrounding membrane environment^3–5^. Emerging structural evidence indicates that, although arrestins engage the active receptor core through multiple interaction modes, they rely on conserved structural elements to recognize the phosphorylated receptor C-terminus and to associate with membranes, underscoring the importance of both phosphorylation barcodes and membrane engagement in arrestin function^6–11^.

Arrestins reside primarily in the cytosol, and pre-association with the membrane greatly facilitates their response to activated receptors^12,13^. Upon receptor activation, arrestin binding to the phosphorylated receptor C-terminus represents an initial step toward high-affinity receptor-and/or membrane-bound states^13,14^. Following recruitment, arrestins undergo clathrin-mediated internalization together with activated receptors, during which they encounter dynamic turnover of distinct phosphoinositide species (PIPs)^15^. The changing lipid environment during trafficking could further modulate arrestin conformation and potentially initiate a second wave of arrestin signaling from intracellular compartments^16–18^. Thus, the interplay between membrane association and receptor engagement is central to arrestin-mediated signaling, yet the mechanistic basis has only begun to be elucidated.

GPCRs can be broadly classified based on their arrestin coupling mode^19^. Class B receptors, such as vasopressin receptor 2 (V_2_R), angiotensin II type 1 receptor, and atypical chemokine receptor 3 (ACKR3), form stable, long-lived complexes with arrestins, supporting sustained arrestin recruitment, endosomal signaling, and regulated trafficking^19–22^. In contrast, Class A receptors, including the β_2_-adrenergic receptor, μ-opioid receptor, and dopamine D_1_ receptor, interact transiently with arrestins, favoring rapid dissociation and receptor recycling^19,23,24^. Whether these distinct arrestin binding dynamics are encoded by receptor phosphorylation barcodes remains an open question. These barcodes are installed by GPCR kinases (GRKs) and vary substantially depending on receptor sequence, GRK isoform, and ligand^5,25–29^.

Prior studies have largely relied on soluble phosphopeptides to examine the effects of barcodes on arrestins^30–32^. Here, we employ N-terminally palmitoylated phosphopeptides tethered to nanodiscs with defined lipid compositions, enabling direct investigation of the interplay between membrane environment and the phosphorylation barcode^33^. Using double electron–electron resonance (DEER) spectroscopy and fluorescence resonance energy transfer (FRET) to track the release of Arr2 autoinhibitory C-tail^33–35^, we find the synergistic impact of lipid bilayer and phosphobarcode in promoting Arr2 activation. By titrating anionic lipids and varying PIP headgroups, we distinguish two modes of membrane control, driven by negative charge, and PIP headgroup identity. Systematic mutagenesis identifies potent phosphopeptide barcode features that support Arr2 activation and membrane engagement. Finally, we explore lipidated phosphopeptides as inhibitors targeting GPCR−Arr2 interactions in live cells and as tools to facilitate structural studies of Arr2−phosphopeptide complexes using cryo-electron microscopy (cryo-EM). Together, these findings establish a multilayered code through which arrestins interpret diverse receptor phosphorylation patterns and lipid cues.

### Membrane Association is Required for Phosphopeptide-mediated Arr2 Activation

The recent cryo-EM structure of ACKR3 in complex with Arr2 and Arr3 reveals that the interface is primarily supported by tail interactions between the N-domain of arrestins and the phosphorylated C-terminus of ACKR3, along with membrane interactions between the arrestins and the surrounding detergent micelles (**Fig. 1A**)^8^. These unique complexes do not require arrestins to engage the activated receptor core, prompting us to test whether core interaction is dispensable and whether the tail and membrane interactions are sufficient to activate and recruit arrestins. To this end, we utilized a lipidated phosphopeptide derived from the unique GRK5 phosphorylation sites located in the proximal region of the ACKR3 C-terminus (**Fig. 1B**)^33^. This peptide, ACKR3_pp1_, contains three phosphorylation sites: pSer335, pThr338 and pThr341, and an N-terminal palmitoylation for tethering to model membrane systems such as nanodiscs. Our recent study shows that in the presence of nanodiscs, ACKR3_pp1_ promotes the release of the auto-inhibitory C-tail of arrestins, rendering it accessible to trypsin digestion, suggesting a synergistic role of membrane association and tail interaction in arrestin activation^33^.

**Figure 1.**
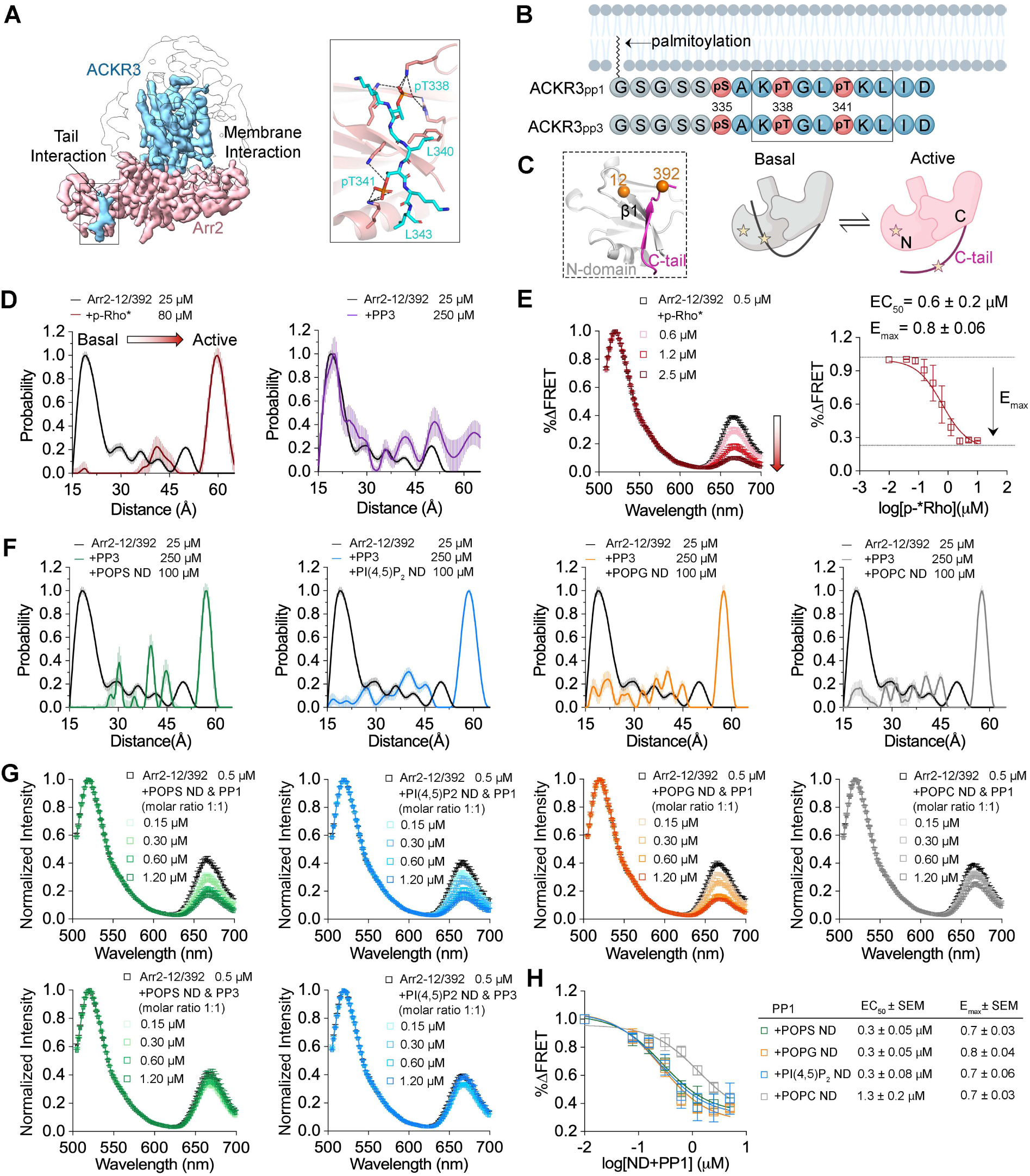
Arr2 activation requires both phosphopeptide binding and membrane association. **A.** Cryo-EM map of ACKR3 phosphorylated by GRK5 in complex with Arr2 (EMD-47700), showing that Arr2 primarily interacts through the phosphorylated receptor C-tail and membrane contacts. The molecular details of the tail interaction are shown (PDB ID 9E82). **B.** Design of ACKR3_pp1_ and ACKR3_pp3_. Both peptides contain three phosphorylation sites (pink spheres). Native sequences derived from ACKR3 are shown as blue spheres, whereas Gly and Ser residues serving as flexible linkers are shown as grey spheres. ACKR3_pp1_ has an N-terminal palmitoylation that facilitates its tethering to nanodiscs. The segment involved in Arr2 binding as revealed in the complex structure (PDB ID 9E82) is highlighted with a rectangular box. **C.** Schematics illustrating that arrestin activation involves release of the auto-inhibitory C-tail from the N-domain. The insert shows the positions of Cα atoms of Ala12 and Ser392 in Arr2 at the basal state (PDB ID 1G4M). Those two sites were labeled for DEER and FRET measurements. **D.** DEER distance distributions of Arr2-12/392 in the absence or the presence of p-Rho* or ACKR3_pp3_. The distance shifts from ∼19Å in the basal state to >55 Å in the presence of p-Rho*, indicating Arr2 activation. These data were previously published^33^. In contrast, ACKR3_pp3_ does not activate Arr2. Error bars represent standard deviation (S.D.). **E.** FRET spectra of Arr2 in the absence or the presence of increasing concentrations of p-Rho*, normalized to the donor peak. The FRET signal decreases in a dose-dependent manner, consistent with Arr2 activation. The FRET signal change is plotted against the p-Rho* concentration to determine the EC_50_ value and E_max_, providing a quantitative measure of Arr2 activation. **F.** The distance distribution shifts from ∼19 Å basal state to longer distances, indicating Arr2 activation in the presence of nanodiscs and ACKR3_pp3_. **G.** FRET spectra of Arr2 in the absence or the presence of increasing concentrations of nanodiscs and ACKR3_pp1_ or ACKR3_pp3_ mixtures, normalized to the donor peak. Nanodiscs in complex with ACKR3_pp1_, but not those with ACKR3_pp3_, cause a dose-dependent decrease in FRET signal, consistent with Arr2 activation. **H.** The FRET signal change is plotted as a function of nanodisc and ACKR3_pp1_ concentrations. The EC_50_ values are comparable among nanodiscs containing anionic lipids and are approximately fourfold lower than that of POPC nanodiscs.

To further characterize the activation process of Arr2, we employed DEER and FRET approaches here. Both methods monitor the release of the auto-inhibitory C-tail of Arr2 as a readout for activation (**Fig. 1C**). We first introduced a pair of cysteine mutations at positions Ala12 and Ser392 into an Arr2 variant in which all native cysteine residues had been mutated while retaining its function^36^. Ala12 is located on the β1-strand of the N-domain, while Ser392 is in the C-tail (**Fig. 1C**). These two cysteines were labeled with nitroxide spin labels for DEER or Alexa Fluor 488 and Atto 647N fluorophores for FRET measurements^30,34^. In the basal state, the C-tail of Arr2 is docked to the N-domain, bringing the two sites into close proximity, approximately 19 Å as measured by DEER (**Fig. 1D**, black curve), and resulting in a robust FRET signal (**Fig. 1E**, black curve). To validate both methods, we used phosphorylated active rhodopsin (p-Rho*), as it is well-established that p-Rho* binds arrestins and displaces the arrestin C-tail from the N-domain^37–39^. As expected, p-Rho* binding led to a shift in the DEER distance to >55 Å (**Fig. 1D**, red curve)^33^ and a dose-dependent reduction in the FRET signal (**Fig. 1E**).

We first employed DEER to test whether a soluble peptide ACKR3_pp3_ (**Fig. 1B**), which contains the same primary protein sequence and phosphorylation sites as ACKR3_pp1_ but lacks the N-terminal palmitoylation, can activate Arr2. Even at a 10-fold molar excess at 250 µM, ACKR3_pp3_ alone did not induce a substantial shift in DEER distance, with the major population remaining at ∼19 Å, indicative of the basal state (**Fig. 1D**). Notably, supplementation with 50 µM nanodiscs, regardless of lipid composition, largely displaces the C-tail, as evidenced by the loss of the 19 Å population and a shift toward longer distances (**Fig. 1F**), indicating C-tail release and Arr2 activation. Nanodiscs used in this study are approximately 9 nm in diameter, a size that we have previously shown to be sufficient for arrestin coupling^33^. Four types of nanodiscs were tested: 100% 1-palmitoyl-2-oleoyl-sn-glycero-3-phosphocholine (POPC), 40%1-palmitoyl-2-oleoyl-sn-glycero-3-phospho-L-serine (POPS) with 60% POPC (molar ratio), 40% 1-palmitoyl-2-oleoyl-sn-glycero-3-phospho-(1’-rac-glycerol) (POPG) with 60% POPC and 10% L-α-phosphatidylinositol-4,5-bisphosphate (PI(4,5)P_2_) with 90% POPC. These results demonstrated that membrane binding plays a key role in phospho-peptide-triggered Arr2 activation.

However, we did not observe any changes in FRET upon the addition of ACKR3_pp3_ and nanodiscs together (**Fig. 1G, Extended Data Fig. 1A**), likely due to the much lower concentrations of Arr2, peptide and nanodiscs used in the FRET assay compared to the DEER experiment. In contrast, ACKR3_pp1_ together with nanodiscs induced a dose-dependent decrease in the FRET signal (**Fig. 1G**). This effect was dependent on phosphorylation, as ACKR3_p4_, which contains the N-terminal palmitoylation but lacks the phosphates, failed to trigger a change in FRET signal (**Extended Data Fig. 1B, C**). It is likely that membrane tethering boosts the local concentration of phosphopeptide near the membrane, thereby promoting effective Arr2 activation. Notably, all nanodiscs containing anionic lipids elicited similar effects, with an EC_50_ of approximately 0.3 μM, significantly more potent than the nanodiscs composed solely of POPC (EC_50_=1.3±0.2 µM) (**Fig. 1H**). These findings further underscore the essential role of anionic lipids in phosphopeptide-induced Arr2 activation.

### Phosphoinositides Regulate Arr2 Activation and Membrane Association

PIPs are signaling lipids with distinct subcellular distributions that are tightly regulated by phosphoinositide kinases and phosphatases^15^. Seven naturally occurring PIP species exist, consisting of mono-, bis-, and tri-phosphorylated forms of phosphatidylinositol (PI) (**Fig. 2A**). Among these, PI(4,5)P_2_ is highly enriched at the plasma membrane and plays a direct role in GPCR signaling, serving as the substrate for G_q_-activated phospholipase^40–42^. Previous studies have thus largely focused on PI(4,5)P_2_^30,43,44^, as Arr2 shows high selectivity for this lipid^33,45^. However, because arrestins encounter dynamic PIP turnovers during internalization and trafficking^46,47^, and because arrestins also contribute to sustained signaling from intracellular compartments enriched in different PIPs^21,48,49^, we sought to examine Arr2 interactions with all seven cellular PIPs.

**Figure 2.**
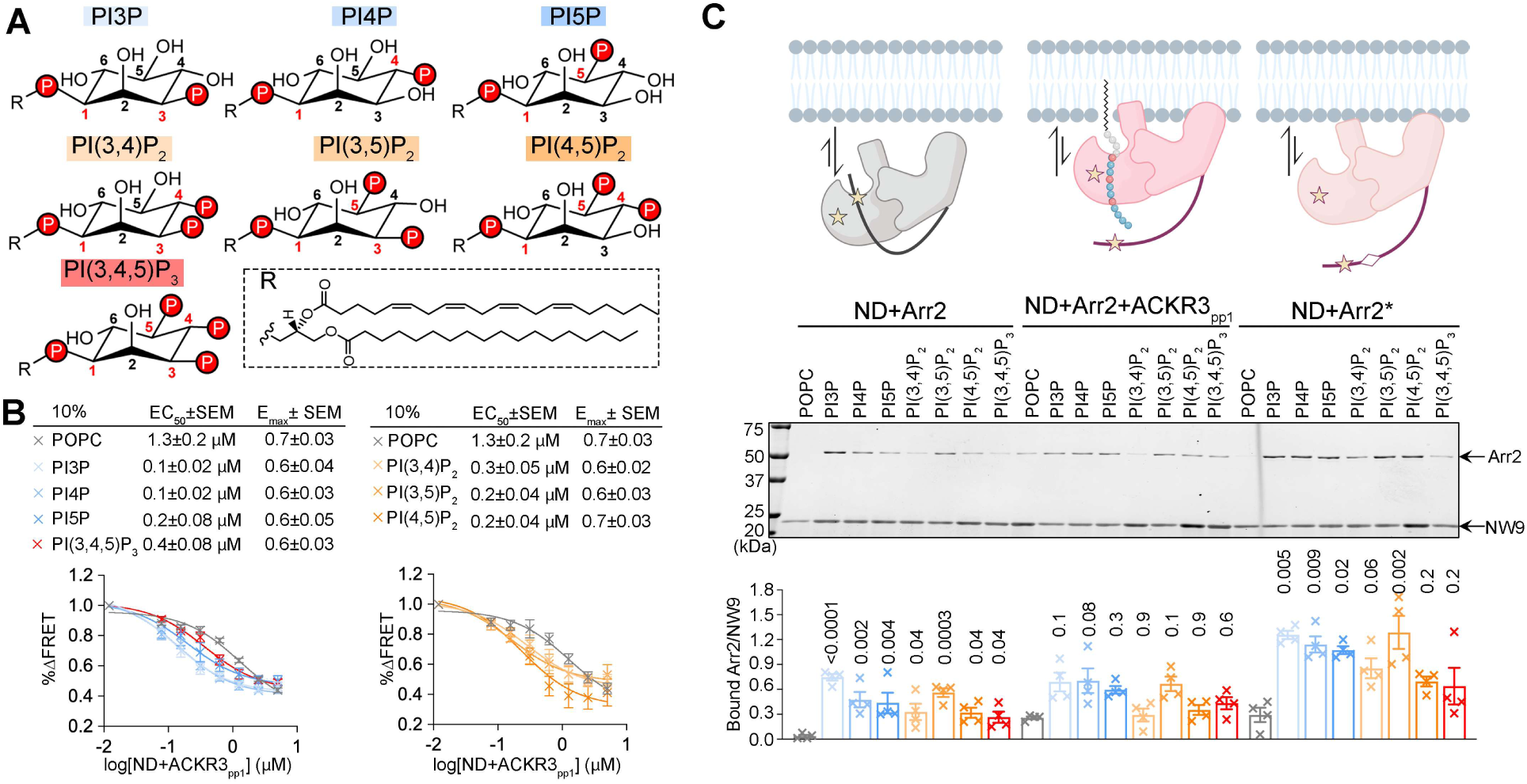
PIPs regulate Arr2 activation and membrane association. **A.** Chemical structures of the PIPs tested in this study. The red spheres highlight the phosphate groups on the inositol ring. **B.** Increasing concentrations of nanodiscs containing 10% PIP and 90%POPC, pre-incubated with an equimolar amount of ACKR3_pp1_, were used to activate Arr2. The normalized decrease in FRET signal was plotted to determine the EC_50_ and E_max_ values. **C.** The scaffolding protein NW9 forms ∼9 nm nanodiscs (ND) and carries a His tag for pull-down assays. Nanodiscs (4 µM) were used in the presence or absence of ACKR3_pp1_ (8 µM) to pull down Arr2 (4 µM) or Arr2* (4 µM). The ratio of bound arrestin band density to NW9 band density was plotted and compared using one-way ANOVA, followed by a Holm–Šídák multiple comparison test, within the ND+Arr2, ND+Arr2+ACKR3_pp1_ and ND+Arr2* groups, respectively. Error bars represent standard error (S.E.) (N=4). The p values comparing arrestin binding to PIP nanodiscs versus POPC nanodiscs within the same group are shown above the corresponding columns.

We employed the FRET-based approach described above (**Fig. 1E**) to measure the effects of different PIPs (**Fig. 2A**) on ACKR3_pp1_-induced Arr2 activation. Nanodiscs containing 10% PIPs and 90% POPC were incubated with ACKR3_pp1_ (molar ratio 1:1) and their effects on Arr2 activation were compared with nanodiscs composed of 100% POPC (**Fig. 2B**). Interestingly, the mono-phosphorylated PIPs, such as PI3P and PI4P, are more effective in potentiating Arr2 activation than the bis- and tri-phosphorylated species PI(3,4)P_2_ or PI(3,4,5)P_3_ (**Fig. 2B**). The position of the phosphate also appears to influence Arr2 activation: PI3P and PI4P are more effective than PI5P, although the difference is subtle (**Fig. 2B**).

We next applied a nanodisc pull-down assay to assess how different PIPs affect Arr2 membrane association in various functional states (**Fig. 2C**). Wildtype (WT) Arr2 binding to nanodiscs reflects its pre-association with the membrane, a step that likely facilitates its transition into high-affinity receptor-bound states. All PIPs significantly enhanced WT Arr2 binding to nanodiscs, with Arr2 showing the highest binding to PI3P nanodiscs, approximately twofold higher than to PI(4,5)P_2_ nanodiscs and roughly threefold higher than to PI(3,4,5)P_3_ (**Fig. 2C**). Among bis-phosphorylated PIPs, PI(3,5)P_2_ nanodiscs bound ∼80% more Arr2 than PI(4,5)P_2_ nanodiscs (**Fig. 2C**).

ACKR3_pp1_ combined with nanodiscs closely mimics the phosphorylated receptor C-tail, and our previous study demonstrated that this tail interaction is sufficient to retain Arr2 at the membrane^33^. Although Arr2 binding to PIP nanodiscs was not significantly different from that of POPC nanodiscs in the presence of ACKR3_pp1_, nanodiscs containing PI3P, PI4P, PI5P or PI(3,5)P_2_ appeared to promote Arr2 membrane association more effectively than those containing other PIPs (**Fig. 2C**). This suggests that Arr2 binding to the phosphorylated receptor is modulated by the surrounding lipid environment.

We next examined how pre-activated Arr2 (Arr2*: I386A, V387A, F388A) interacts with nanodiscs containing different PIPs. By disrupting autoinhibitory intramolecular interaction, these triple alanine mutations shift the conformational equilibrium of Arr2 toward intermediate activated states^50^. Arr2* binding to nanodiscs can reflect the functional state in which Arr2* has dissociated from the receptor but remains active. All PIP-containing nanodiscs showed approximately twofold higher binding to Arr2* than to WT Arr2, consistent with our previous finding that Arr2 activation promotes its membrane association^33^. Interestingly, Arr2* displayed a binding pattern similar to that of Arr2 in the presence of ACKR3_pp1_, with higher binding observed for PI3P, PI4P, PI5P and PI(3,5)P_2_ nanodiscs (**Fig. 2C**).

Overall, these results suggest that PIPs promote Arr2 membrane association and activation in the presence of a phosphorylated receptor tail. Mono-phosphorylated species, particularly endosome-enriched PI3P, show the strongest effects, whereas more highly phosphorylated PIP headgroups, such as PI(3,4,5)P_3_, contribute the least.

### Anionic Lipids Potentiate Phosphopeptide-Triggered Arrestin Activation

Anionic lipids potentiate phosphopeptide-triggered Arr2 activation (**Fig. 1H**), whereas for PIPs, additional phosphates appear to reduce their effectiveness in promoting Arr2 activation (**Fig. 2B**). This suggests that negatively charged lipids, such as POPS and POPG, likely promote Arr2 activation through mechanisms distinct from those of PIPs.

To further investigate this, we examined how varying concentrations of POPS influence Arr2 activation and membrane binding (**Fig. 3**). Increasing the POPS content in the nanodiscs led to a clear potentiation of Arr2 activation by ACKR3_pp1_, with approximately an order-of-magnitude increase in potency at 80% POPS (EC_50_ = 0.1±0.04 µM) compared to 0% POPS (EC_50_ = 1.3±0.2 µM) (**Fig. 3A**). We next tested whether the enhanced activation correlates with increased nanodisc binding using pull-down assays (**Fig. 3B**). Significantly enhanced Arr2 nanodisc binding was observed when the POPS concentration exceeded 80% (**Fig. 3B**). These results indicate that Arr2 preferentially interacts with negatively charged membrane surfaces, which synergistically potentiate its activation in the presence of a phosphorylated receptor C-tail (**Fig. 3A**). Consistent with our prior findings, Arr2* exhibited overall higher nanodisc binding than Arr2 and showed a clear POPS-dependent increase in membrane association (**Fig. 3B**).

**Figure 3.**
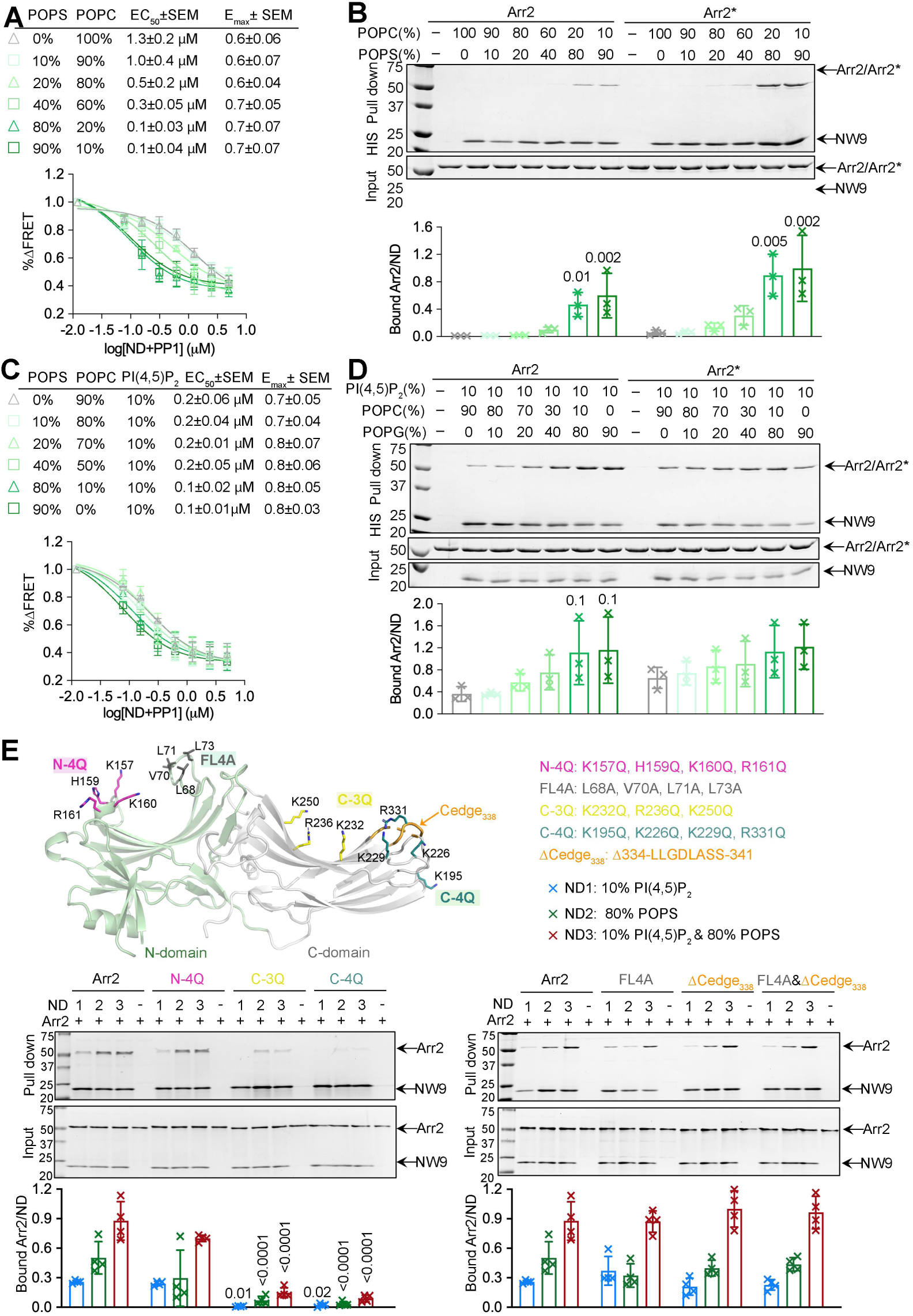
Anionic phospholipids potentiate phosphopeptide-induced Arr2 activation and membrane association. **A.** Nanodiscs containing increasing concentrations of POPS, pre-incubated with an equimolar amount of ACKR3_pp1_, were used to activate Arr2. **B.** Nanodiscs containing increasing concentrations of POPS (4 µM) were used to pull down Arr2 (5 µM) or Arr2* (5 µM). **C.** Nanodiscs containing 10% PI(4,5)P_2_ and increasing concentrations of POPS, pre-incubated with an equimolar amount of ACKR3_pp1_, were used to activate Arr2. **D.** Nanodiscs containing 10% PI(4,5)P_2_ and increasing concentrations of POPS (4 µM) were used to pull down Arr2 (5 µM) or Arr2* (5 µM). **E.** The structure of Arr2 in the basal state (PDB ID 1G4M) highlights the positions of the mutants tested in the pull-down assays. Nanodiscs containing 10%PI(4,5)P_2_ (ND1), 80%POPS (ND2) or 80% POPS&10%PI(4,5)P_2_ (ND3) (4 µM) were used to pull down Arr2 WT and variants (5 µM). For panels **A, C,** the normalized decrease in FRET signal was plotted to determine the EC_50_ and E_max_ values. For panels **B, D, E,** the ratio of bound arrestin band density to NW9 band density was quantified. Error bars represent standard deviation (S.D.) (N=3 for B, D; N=4 for E). For panels **B** and **D**, the ratio of bound Arr2 to nanodisc was compared using one-way ANOVA, followed by a Holm–Šídák multiple comparison test, within the Arr2 and Arr2* groups, respectively. p values ≤ 0.05 for comparisons of arrestin binding to POPC nanodiscs within the same group are shown above the corresponding columns. **E.** The ratio of bound Arr2 to nanodisc was compared using two-way ANOVA followed by a Dunnett’s multiple comparison test, within the ND1, ND2 and ND3 groups, respectively. p values ≤ 0.05 for comparisons of Arr2 WT to variants within the same nanodisc condition are shown above the corresponding columns.

To test whether the effects of POPS and PI(4,5)P_2_ are additive, we next performed the same POPS titration in the presence of 10% PI(4,5)P_2_. As shown above, 10% PI(4,5)P_2_ greatly facilitates Arr2 activation (**Fig. 1H**); nevertheless, increasing the POPS content up to 80% further enhanced Arr2 activation, lowering the EC_50_ to 0.1 µM (**Fig. 3C**). Notably, with nanodiscs containing high POPS content (80%-90%), PI(4,5)P_2_ does not make any difference in ACKR3_pp1_-induced Arr2 activation anymore (**Fig. 3A, C**). POPS further promotes Arr2 binding to PI(4,5)P_2_ nanodiscs in a dose-dependent manner, indicating that these two lipids can cooperatively retain Arr2 at the membrane (**Fig. 3D**).

Given the additive effects observed in Arr2 activation and membrane binding (**Fig. 3A-D**), we reasoned that multiple structural elements are likely involved in coordinating binding to PI(4,5)P_2_ and POPS. To investigate this, we purified a series of Arr2 variants in which clusters of Lys and Arg residues on the N- and C-domains were substituted with Gln, including the canonical (C-3Q)^45^ and non-canonical PI(4,5)P_2_ (C-4Q)^43^ binding sites (**Fig. 3E**). Pull-down assays revealed that both the C-3Q and C-4Q variants exhibited >90% reduction in binding to all tested nanodiscs compared to WT Arr2, highlighting the importance of these C-domain residues in lipid binding. In contrast, the N-4Q variant showed comparable binding to POPS and PI(4,5)P_2_ nanodiscs.

We next mutated the hydrophobic residues on the finger loop (FL4A) and/or deleted part of the C-edge (ΔCedge_338_) to assess whether the hydrophobic anchors contribute to nanodisc binding. Neither mutation resulted in significant differences in binding to POPS or PI(4,5)P_2_ nanodiscs compared to WT Arr2 (**Fig. 3E**). All Arr2 variants were correctly folded and functional, as confirmed by trypsin digestion of the autoinhibitory C-terminus upon binding to p-Rho*(**Extended Data Fig. 2**). Collectively, these results indicate that multiple sites on the C-domain cooperate to mediate binding to PI(4,5)P_2_ and other negatively charged lipids such as POPS. Because the phosphorylated receptor C-terminus engages Arr2 on the N-domain, this interaction likely generates a force that facilitates interdomain rotation and activation.

### Defining Phosphorylation Barcodes that Drive Arr2 Activation

We next sought to understand how specific features of the phosphorylation barcode influence arrestin activation. Because phosphates added at proximal sites by GRK5 or distal positions by GRK2 on ACKR3 (**Fig. 4A**) can alter the configuration of the receptor–arrestin complex^8^, we first examined whether the distance between the phosphorylation site and the membrane affects Arr2 activation (**Fig. 4A**). To this end, we designed ACKR3_pp2_, which is identical to ACKR3_pp1_ except that it contains 11 additional residues between the palmitoylation and the phosphorylation sites (**Fig. 4A**). Although the EC_50_ values are comparable, the E_max_ of ACKR3_pp2_ is ∼25% lower than that of ACKR3_pp1_, indicating that the position of the barcode relative to the membrane also plays a role in Arr2 activation. In the extreme case, ACKR3_pp3_, which is not tethered to the membrane, showed no detectable activation within the tested concentration range (**Fig. 1F**), suggesting that the greater the distance of the barcode from the receptor core, the less efficient it is in activating Arr2.

**Figure 4.**
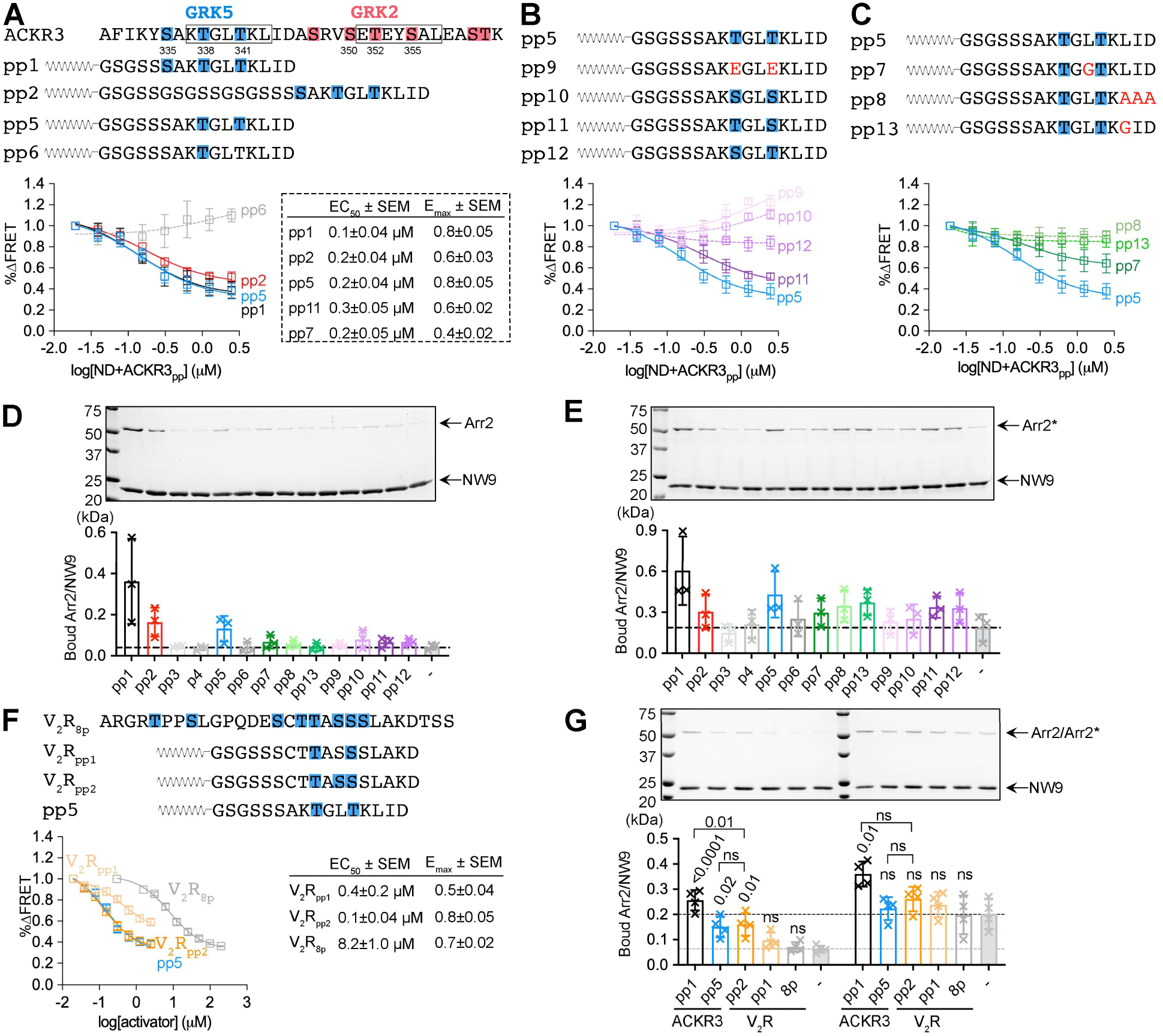
Barcode features that regulate Arr2 activation and membrane association. **A-C**. The list of designed ACKR3 peptides and the corresponding FRET curves reflecting their potency in Arr2 activation is shown. The phosphorylation sites are highlighted in blue. Increasing concentrations of nanodiscs containing 10%PI(4,5)P_2_, 40%POPG, and 50%POPC, pre-incubated with a twofold molar excess of peptide, were used to activate Arr2. The normalized decrease in FRET signal was plotted to determine the EC_50_ and E_max_ values. **D, E.** Nanodiscs (4 µM) containing 20% POPS and 80% POPC, pre-incubated with each peptide (8 µM) were used to pull down Arr2 (5 µM) (**D**) or Arr2* (5 µM) (**E**). The ratio of bound Arr2 (**D**) or Arr2* (**E**) to NW9 normalized to ACKR3_pp1_ was plotted. **F.** The sequence alignment of V_2_R_8p_, V_2_R_pp1_, V_2_R_pp2_ and ACKR3_pp5_ and their corresponding FRET curves reflecting their potency in Arr2 activation. The phosphorylation sites are highlighted in blue. **G.** Nanodiscs (4 µM) containing 20% POPS and 80% POPC, pre-incubated with each peptide (8 µM) were used to pull down Arr2 (5 µM) or Arr2* (5 µM). The ratio of bound Arr2 or Arr2* to NW9 normalized to ACKR3_pp1_ was plotted.

To determine how many phosphates are required for effective Arr2 activation, we synthesized lipidated peptides containing either two (ACKR3_pp5_) or one phosphate group (ACKR3_pp6_) (**Fig. 4A**). ACKR3_pp5_ effectively activated Arr2 with EC_50_ and E_max_ values comparable to those of ACKR3_pp1_, whereas ACKR3_pp6_ showed minimal activity. These results suggest that two phosphates are both necessary and sufficient for robust Arr2 activation.

To examine whether the chemical identity of the phosphosites affects Arr2 activation, we substituted the pThr with either Glu (ACKR3_pp9_) or pSer (ACKR3_pp10_) (**Fig. 4B**). Glu residues were ineffective at activating Arr2; unexpectedly, pSer also failed to induce robust Arr2 activation (**Fig. 4B**), despite containing the same phosphate moiety as pThr. This observation suggests that the additional methyl group on pThr likely contributes to specific structural or conformational features required for optimal Arr2 activation. To further test this idea, we next replaced each pThr with pSer individually (ACKR3_pp11_ and ACKR3_pp12_) (**Fig. 4B**). ACKR3_pp11_ activated Arr2 more effectively than ACKR3_pp12_, but remained less potent than ACKR3_pp5_, suggesting that both pThr residues enhance Arr2 activation, with the first site contributing more than the second.

In addition to the electrostatic interactions between the phosphates on the receptor C-terminus and Lys/Arg residues on the Arr2 N-domain, we also observed hydrophobic contacts that may contribute to receptor−arrestin interactions^8^ (**Fig. 1A**). To test this, we synthesized peptides carrying the same phosphorylation pattern as ACKR3_pp5_ but with specific residue substitutions in the flanking sequences: Leu340 was mutated to Gly (ACKR3_pp7_) and Leu343-Asp345 was replaced with a triple Ala sequence (ACKR3_pp8_) (**Fig. 4C**). ACKR3_pp7_ exhibited an EC_50_ value comparable to that of ACKR3_pp5_ but showed a ∼50% reduced E_max_, whereas ACKR3_pp8_ barely activated Arr2 (**Fig. 4C**). Since our recent structure suggests that Leu343 directly interacts with the Arr2 N-domain, we further mutated it to Gly (ACKR3_pp13_). This single-residue substitution produced effects similar to ACKR3_pp8_ (**Fig. 4C**), further emphasizing that the residues flanking the phosphorylation sites play a crucial role in Arr2 activation.

### Define the Phosphorylation Barcode that Promotes Arr2 Membrane Association

We next examined how these peptides influence membrane recruitment of Arr2. Nanodiscs containing 20% POPS and 80% POPC were used to pull down Arr2 in the presence of various peptides (**Fig. 4D**). Because the binding site of the phosphorylated receptor C-terminal peptide overlaps with that of the Arr2 autoinhibitory C-tail, these elements compete for access to the Arr2 N-domain. Thus, it is reasonable that the most potent barcodes for Arr2 activation also promote the strongest membrane association of Arr2. Indeed, ACKR3_pp1_ promoted the highest Arr2 binding to the nanodiscs (**Fig. 4D**). Notably, ACKR3_pp1_ induced approximately two-fold higher Arr2 binding compared to ACKR3_pp2_, indicating that the spacer between the transmembrane (TM) domain and the barcode plays an important role in Arr2 membrane recruitment (**Fig. 4D**). Although ACKR3_pp1_ and ACKR3_pp5_ showed comparable potency in Arr2 activation (**Fig. 4A**), ACKR3_pp1_ further enhanced Arr2 membrane association (**Fig. 4D**), likely due to the additional phosphate increasing local negative charge, analogous to the potentiating effect of negatively charged lipids (**Fig. 3B**).

It is striking that all other barcodes failed to facilitate Arr2 membrane association under the tested conditions (**Fig. 4D**). We next examined whether these peptides influence the membrane association of Arr2*. By partially bypassing the activation process, ACKR3_pp8_ and ACKR3_pp13_ promoted Arr2* membrane binding to an extent comparable to ACKR3_pp5_, indicating that residues following the phosphorylation sites are less important for membrane association (**Fig. 4E**) than for triggering Arr2 activation (**Fig. 4C**). Additionally, pThr residues were critical for retaining Arr2* at the membrane, as substitution of both with pSer or Glu residues markedly reduced membrane association (**Fig. 4E**). Finally, extension of the spacer length, as in ACKR3_pp2_, resulted in an approximately two-fold reduction in Arr2* binding relative to ACKR3_pp1_ (**Fig. 4E**).

Altogether, this allowed us to rank barcodes based on their potency in activating Arr2 and promoting Arr2 membrane association, with the most effective motif being pTxΦpTxΦ (x, any residue; Φ, hydrophobic residue), followed by pTxΦpSxΦ and pTxxpS/TxΦ. Importantly, the position of the barcode relative to the receptor TM core or the membrane is also critical, supporting the concept of a regional barcode.

### Engineering V_2_R_pp_ for Potent Arr2 Activation and Robust Membrane Association

Because the barcodes tested thus far were derived from ACKR3, which is naturally biased toward arrestins, we next sought to determine whether this membrane-mediated potentiation of Arr2 activation also applies to canonical GPCRs. To this end, we examined the effects of a widely used hyperphosphorylated peptide derived from vasopressin receptor 2 (V_2_R_8p_)^51^. V_2_R_8p_, which contains eight phosphates within a single peptide, could trigger Arr2 activation, albeit with a much higher EC_50_ of ∼8 μM as measured by FRET (**Fig. 4F**). When applied at high concentration (200 μM), the major population of Arr2-12/392 displayed a shift in DEER distance to >55 Å (**Extended Data Fig. 3**), indicative of activation.

V_2_R_8p_ contains a barcode of pTxpSpSxΦ, similar to the ACKR3 barcode pTxxpS/TxΦ. We therefore designed two peptides, V_2_R_pp1_ and V_2_R_pp2_, containing two or three phosphates, respectively, each with an N-terminal palmitoylation to promote membrane association (**Fig. 4F**). In the presence of nanodiscs, the lipidated V_2_R_pp1_ and V_2_R_pp2_ were potent activators of Arr2. V_2_R_pp2_ showed an approximately 80-fold decrease in EC_50_ with a comparable E_max_ relative to V_2_R_8p_, indicating that membrane association plays a far more important role than the number of phosphates. V_2_R_pp1_ was slightly less efficient than V_2_R_pp2_ in activating Arr2 (**Fig. 4F**), indicating that the third position in the barcode could be either a phosphate (V_2_R_pp2_) or a hydrophobic residue (ACKR3_pp5_) for robust Arr2 activation. Indeed, the pTxpSpSxΦ motif falls within the broad pS/TxpS/TpS/T (“PxPP”) barcode class proposed by Shukla and colleagues^52^.

We further tested how these peptides impact Arr2 and Arr2* membrane binding. V_2_R_pp2_ promoted significantly higher Arr2 membrane binding than V_2_R_pp1_, at a level comparable to ACKR3_pp5_ but lower than that of ACKR3_pp1_ (**Fig. 4G**). Arr2* exhibited enhanced membrane binding compared to Arr2 and the addition of V_2_R_pp1_ or V_2_R_pp2_ did not further promote Arr2* membrane binding (**Fig. 4G**), as with ACKR3_pp5_. Altogether, our results strongly suggest that membrane-mediated potentiation of Arr2 activation is broadly applicable and together with the phosphorylation barcode, can determine the efficiency in initiating arrestin-mediated responses.

### Global Analysis of Regional Phosphorylation Motifs Within the GPCR Family

Based on their potency in triggering Arr2 activation and promoting membrane association, we ranked the phosphorylation barcodes, with the most effective motif being pTxΦpTxΦ, followed by pTxΦpSxΦ, pTxxpS/TxΦ and pS/pTxxpS/TxΦ (Motif I-IV; Φ=[VILMFYW]; x=any residue). We then performed a proteome-wide search to identify these motifs in human GPCR sequences, mapped their positions relative to TM7, and curated available experimental evidence for arrestin coupling to classify receptors as transient (Class A) or sustained binders (Class B), or arrestin-uncoupled (**Fig. 5; Extended Data Fig. 4A, Supplementary Table 1**).

**Figure 5.**
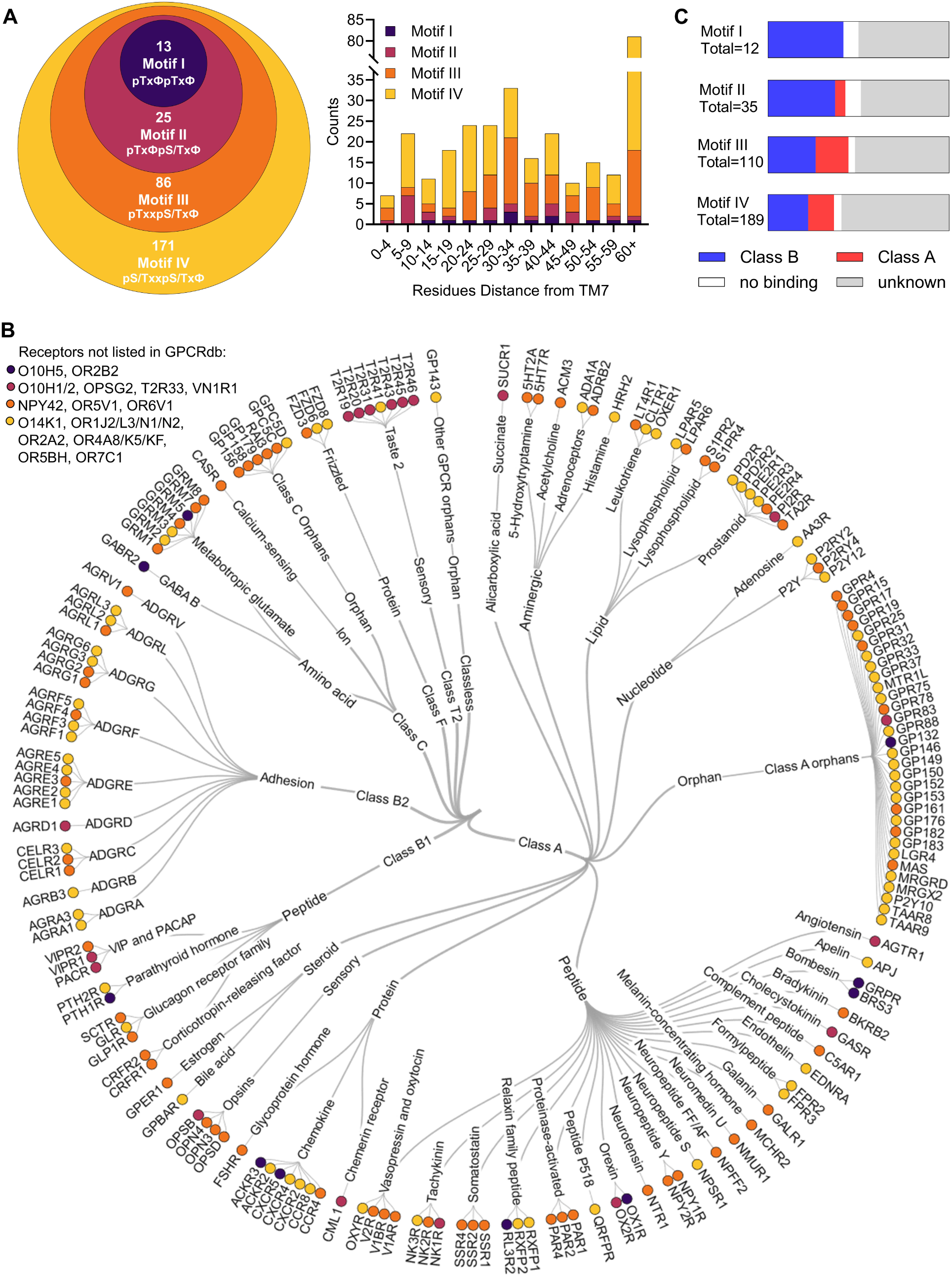
Global analysis of phosphorylation motifs across the human GPCRome. **A.** ACKR3-derived phosphorylation motifs (Motif I-IV) identified across human GPCR sequences, together with their location distribution relative to TM7. **B.** Phylogenetic tree of GPCRs carrying Motif I–IV, with nodes colored according to the highest-potency motif detected in each receptor (see panel A). Receptors not annotated in GPCRdb are listed separately. **C.** Distribution of arrestin coupling classes: sustained (Class B), transient (Class A), no detectable arrestin binding, or unknown, among receptors carrying each phosphorylation motif. See Supplemental Table 1 for motif occurrences and curated arrestin-coupling assignments.

Motif I was rare across the GPCRome, occurring at only 13 sites across 12 receptors, including ACKR3 and several peptide hormone receptors known to form stable arrestin complexes, such as parathyroid hormone 1 receptor (PTH_1_R), orexin receptor type 1 (OX_1_R) and gastrin-releasing peptide receptor (GRPR)^53–55^. Motifs II-IV were progressively more prevalent and broadly distributed across receptor families, spanning both well-characterized arrestin-binding GPCRs and underexplored groups such as adhesion GPCRs, metabotropic glutamate receptors, and numerous orphan and olfactory GPCRs. Notably, a substantial fraction of identified receptors are involved in sensory perception, including light, taste and olfaction. For example, a cluster of bitter taste receptors (Tas2Rs) carries a conserved Motif II within the proximal C-terminus (**Fig. 5B, Extended Data Fig. 4B**).

Among receptors with available experimental data on arrestin coupling, Class B behavior was more frequently observed in receptors containing Motif I (83%) or Motif II (72%) than in those containing Motif III (55%) or Motif IV (55%) (**Fig. 5C, Supplementary Table 1**). This trend is consistent with a positive relationship between stronger barcode potency and sustained arrestin coupling. Across all motifs, 73% of sites were located within 60 residues of the TM domain, with a peak at 25-34 residues downstream of TM7 (**Fig. 5C**). Assuming an average helix 8 length of ∼13 residues^56,57^, this places the barcode on average ∼12-21 residues from the membrane. Notably, the locations of the four barcode motifs span a wide range of distances, from a few residues up to ∼1700 residues in the orphan GPR179 (**Fig. 5A**).

We applied the same analysis to previously described arrestin-binding motifs, including the pS/TxpS/TpS/T (“PxPP”) motif^52,58–60^ and the pS/Tx(x)pS/TxxpS/T (“Rho”) motif^39^ (**Extended Data Fig. 4C, Supplementary Table 1**). These two barcode motifs were primarily defined based on phosphorylation patterns observed in complex structures^39,58^. In contrast, our analysis incorporates the effects of nearby hydrophobic residues and distinguishes between pSer and pThr, informed by both structural insights^8^ and quantitative functional characterization (**Fig. 4**). Motif IV (pS/pTxxpS/TxΦ), which imposes the least sequence restriction, was found at more than 300 sites and substantially overlapped with the PXPP and Rho motifs, consistent with the broad substrate specificity of the non-visual arrestins. Additionally, all motifs exhibit considerable variation in their distance from TM7, potentially fine-tuning arrestin recruitment and diversifying downstream signaling outcomes (**Extended Data Fig. 4B**).

### Lipidated Phosphopeptides As Tools to Tune Arr2−Receptor Interaction in Cells

Since membrane-tethered phosphopeptides are sufficient to activate Arr2 and drive its membrane translocation (**Fig. 4**), we reasoned that they could be used as tools to sequester active Arr2 and thereby reduce Arr2-mediated signal desensitization. This strategy can be particularly valuable for therapeutic interventions targeting receptors that undergo rapid and excessive arrestin-mediated internalization and degradation^1,61–63^. By prolonging receptor residence at the plasma membrane, such drugs could promote more sustained therapeutic effects.

To test this idea, we employed a NanoBiT-based arrestin-recruitment assay^64^ to compare Arr2−receptor coupling in the presence or absence of various peptides (**Fig. 6A, B**). SmBiT and LgBiT were fused to the receptor C-terminus and the Arr2 N-terminus, respectively. Upon agonist stimulation, Arr2 recruitment brings the two luciferase fragments into close proximity, generating a luminescent signal. A lipid-tethered phosphopeptide can potentially compete for Arr2 binding and thereby dampen the luminescence (**Fig. 6A**)^65^. We therefore compared agonist-induced luminescence in cells with or without peptide pre-treatment to quantify the inhibitory effect of each peptide (**Fig. 6C, D**).

**Figure 6:**
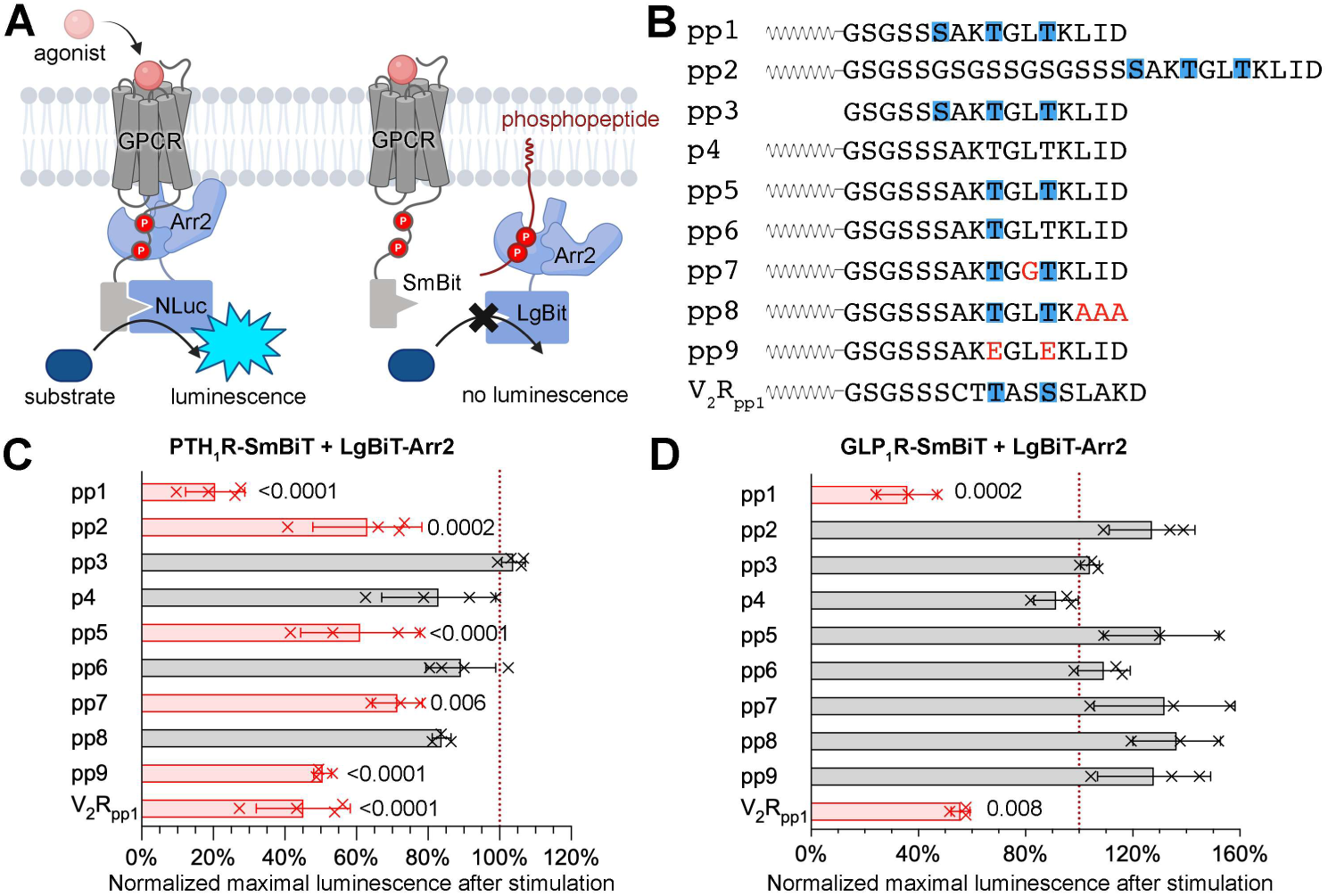
Lipidated phosphopeptides modulate GPCR−Arr2 interactions in live cells. **A.** Schematic illustrating the NanoBiT assay for assessing GPCR−Arr2 interaction. SmBiT and LgBiT were fused to the receptor C-terminus and the Arr2 N-terminus, respectively. Upon agonist stimulation, Arr2 is recruited to the activated receptor, bringing SmBiT and LgBiT into close proximity, generating a luminescent signal. A lipidated phosphopeptide can sequester Arr2 and prevent its association with the receptor, thereby reducing luminescence. Created in BioRender. **B.** List of tested peptides. Blue boxes indicate phosphorylation sites, and the mutations are highlighted in red. pp1-pp9 are derived from ACKR3 and all peptides except pp3 contain a N-terminal palmitoylation site. **C** and **D.** HEK293 cells were pre-incubated with peptide or vehicle; luminescence after agonist stimulation was recorded for PTH_1_R (**C**) or GLP_1_R (**D**). The graphs show maximum luminescence from peptide-treated cells normalized to the vehicle control. Normalized values were compared using one-way ANOVA, followed by a Tukey multiple comparison test against the pp3 treatment group. Error bars represent standard deviation (S.D.) from at least three biological replicates. The groups with p values < 0.05 for comparisons against the pp3 treatment group are highlighted in pink with the corresponding p values shown next to each column.

We selected two secretin-like GPCRs, GLP_1_R and PTH_1_R, as both robustly recruit Arr2^53,66,67^. ACKR3_pp1_, carrying the most potent barcode for promoting Arr2 activation and membrane association in vitro (**Fig. 4A, E**), also exhibited the strongest inhibition in cells, reducing luminescence signals of PTH_1_R−Arr2 and GLP_1_R−Arr2 by ∼80% and ∼65%, respectively (**Fig. 6C, D**). Notably, different peptides showed varying levels of inhibition of Arr2 association with PTH_1_R (**Fig. 6C**), suggesting that phosphopeptide design can be tailored to fine-tune Arr2 signaling. In contrast, most peptides were ineffective at blocking Arr2 interaction with GLP_1_R (**Fig. 6D**), showing that inhibition is receptor-dependent, potentially reflecting stronger tail or core interactions that stabilize the GLP_1_R−Arr2 complex in this cellular context.

Together, these results demonstrate the potential of lipidated phosphopeptides as tools to modulate Arr2-receptor interactions in cells. However, their practical application will require improved delivery strategies, as high micromolar concentrations were necessary to achieve robust inhibition in the NanoBiT assay.

### A generalizable strategy to determine Arr2-phosphopeptide structures

The interactions between phosphorylation barcodes and arrestins have been largely informed by structural insights^5,8,39,58^; however, the high conformational flexibility of active arrestins and the paucity of suitable structural tools have restricted our understanding to only a handful of phosphorylation patterns. We recently developed Fab7, a tool that selectively stabilizes arrestins in the activated states by engaging the arrestin inter-domain region^8^. Fab7 does not interact directly with the receptor or the phosphates and has been successfully used for structural determination^8^. We therefore sought to develop a generalizable strategy to determine structures of Arr2 bound to different phosphopeptides.

Our first attempt at assembling the complex using ACKR3_pp3_, pre-activated Arr2* and Fab7 failed to generate any 2D averages showing defined structural features (**Fig. 7A**). We reasoned that the activated Arr2* could gain substantial conformational flexibility on the receptor- or membrane-binding side on which Fab7 would not have any stabilizing effect. We therefore added nanodiscs and combined the lipidated phosphopeptides with Fab7, which successfully led to defined 2D averages and a 3D reconstruction (**Fig. 7A, B, Extended Data Fig. 5**). Interestingly, the nanodisc density was not observed either in the 2D averages or 3D reconstruction, likely because the nanodiscs are heterogeneous in size and the binding between lipids and proteins is not well-defined. Indeed, the density for the membrane-binding interface on Arr2 is of poor quality and we could not model Arr2 with confidence. Notably, the phospho-peptide shows defined density (**Fig. 7B**), similar to the tail interaction observed in the cryo-EM structure of Arr2 in complex with GRK5-phosphorylated ACKR3^8^ (**Fig. 1A**). Fab7 bound the inter-domain region of Arr2, similarly to what was observed before^8^. This strategy can potentially be generalized to visualize the binding of any phosphopeptide to arrestins, provided that the lipidated peptide can retain Arr2 near the nanodiscs. Although the resolution will likely be limited at the membrane-binding interface, the peptide-binding interface could provide novel structural insights. The nanodiscs could also be a useful tool to stabilize activated arrestins to facilitate the structural determination of arrestin in complex with downstream effectors, for which no high-resolution structural information is currently available.

**Figure 7.**
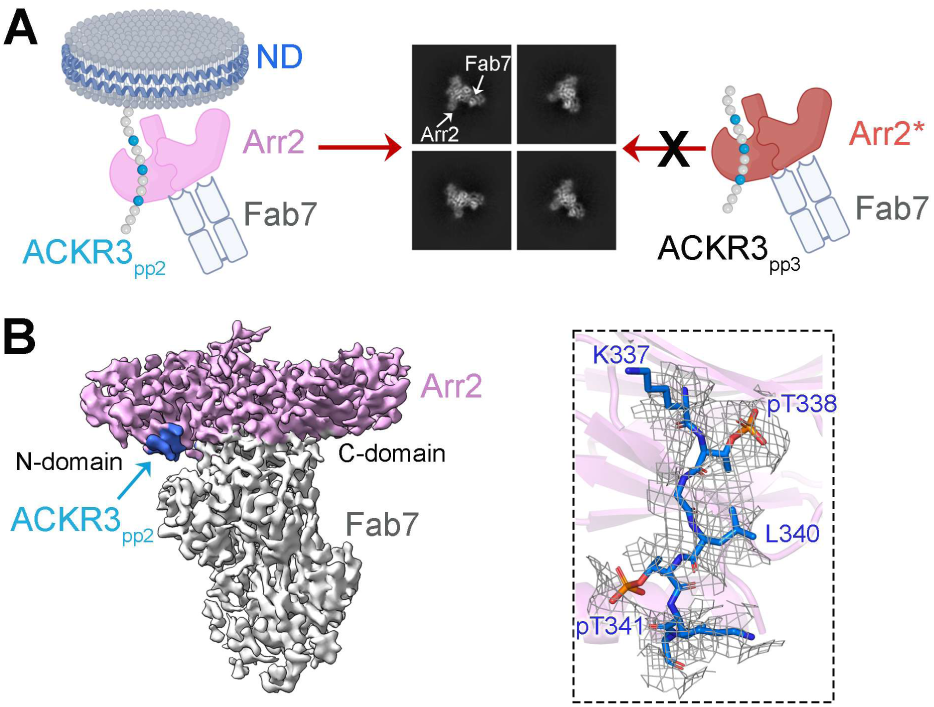
Development of a strategy for determineing Arr2-phosphopeptide structures. **A.** Cartoon illustrating two strategies for assembing the complex for cryo-EM studies. Using the nanodiscs, the lipidated phosphopeptide ACKR3_pp2_, Arr2 and Fab7, but not the soluble phosphopeptide ACKR3_pp3_, Arr2* and Fab7, led to 2D averages with defined features. The density for nanodiscs was not observed in the 2D average. **B.** Cryo-EM reconstruction of Arr2 (pink) in complex with ACKR3_pp2_ (blue) and Fab7 (gray). The electron density and model of ACKR3_pp2_ is shown in the rectanglar.

## Discussion

Arrestins function as molecular coincidence detectors that integrate multiple receptor-derived signals before engaging productively with GPCRs^68^. Our study further extends this concept by highlighting the role of the membrane in this process. Membrane association greatly potentiates Arr2 activation in the presence of a phosphorylated receptor C-tail, likely facilitating its transition to high-affinity receptor-binding or membrane-binding states. This mechanism provides an additional means to fine-tune arrestin-mediated signaling. The same barcode located at different positions along the receptor C-tail would likely produce different magnitudes of arrestin response^8,33^. The GPCR C-termini are highly variable in length, ranging from a few to more than 1700 residues (GPR179). Palmitoylation, clusters of Arg/Lys residues, and/or disorder or absence of H8 can further modulate the distance between the phosphorylation barcode and the membrane, thereby diversifying the arrestin signaling output^69–72^. In addition, the distance from the membrane enables barcode modulation by different GRK subfamilies and other kinases such as PKA, as membrane-proximal sites are preferentially targeted by GRK5/6 and PKA^8,73–75^. This introduces an additional layer of barcoding regulation, since GRK expression levels vary across tissues and pathological conditions, potentially tailoring arrestin-mediated signaling in a context-dependent manner^29^.

ACKR3, a naturally arrestin-biased receptor, contains two clusters of potent phosphorylation barcodes and can effectively couple to arrestins without the involvement of the activated receptor core^8^. By studying phosphopeptides derived from ACKR3, we systematically ranked these barcodes based on their potency in promoting Arr2 activation. Our results indicate that two phosphates are necessary and sufficient for Arr2 activation, with a preference for pThr over pSer or Glu substitutions; neighboring hydrophobic residues also play important roles. Based on these findings, we defined four patterns of barcodes (Motif I−IV) with decreasing potency in Arr2 activation. Indeed, a search for these motifs across the human GPCRome reveals a correlation between high-potency barcodes and sustained (“Class B”) arrestin coupling. Since arrestin coupling relies on GRK phosphorylation, we speculate that these barcodes reflect not only a preference for Arr2 binding but also preferred substrate motifs for GRKs.

Distinct cellular compartments harbor unique PIP species and lipid compositions, functioning as molecular “zipcodes” that direct the trafficking of signaling proteins. We demonstrate that different PIP species exert distinct effects on Arr2 activation and membrane association (**Fig. 2**). Importantly, PI3P, which is highly enriched in intracellular membranes such as endocytic vesicles^46,47^, is the most effective. These findings reveal that Arr2 discriminates among PIP species, establishing a molecular framework for spatially compartmentalized arrestin signaling^76–80^.

Other anionic phospholipids such as POPS are abundant in the plasma membrane, accounting for 40-60% of the inner leaflet^81,82^, where arrestins interact with receptors. We show that these negatively charged lipids also potentiate Arr2 activation and enhance Arr2 membrane binding, either alone or synergistically with PIPs (**Fig. 3A, E**). These findings further highlight that the membrane interaction plays a critical role in arrestin responses. We further demonstrate that the C-domain of Arr2 harbors multiple clusters of Lys and Arg residues, including the canonical and non-canonical PI(4,5)P_2_ binding sites in the C-domain^43,45^, that are responsible for both PI(4,5)P_2_ and POPS binding. Since the phosphorylated receptor C-tail primarily engages Arr2 through the N-domain while the membrane is engaged through the C-domain, this dual-anchoring may generate a mechanical force that promotes domain rotation and thereby Arr2 activation. Given that Arr3’s higher flexibility arises from sequence variation within its C-domain, differences in its membrane interface may contribute to the distinct behavior of the two non-visual arrestins^33,83^. Extending this experimental approach to Arr3 may reveal arrestin subtype-specific weighting of phospho-barcode versus lipid zipcode inputs.

We demonstrated that lipidated phosphopeptides inhibit agonist-dependent arrestin recruitment to PTH_1_R and GLP_1_R in cells. A recent study showed that a lipidated peptide derived from V_2_R with Glu residue substitutions for phosphorylation potentiated morphine’s analgesic effect in mice^65^. Lipidated peptides represent promising tools that could be tailored to inhibit arrestin recruitment to specific GPCRs for therapeutic purposes. However, our study highlights a dilemma for peptide design: Glu substitutions lead to a marked reduction in Arr2 activation potency, whereas phosphorylation limits permeability. Effective translation of peptide tools into robust cellular modulators will thus require high-potency designs combined with enhanced stability and cellular delivery.

By extending this approach to a classic GPCR C-terminus (V_2_R_8p_) (**Fig. 4F**), we demonstrated that the combination of lipidated phosphopeptides and nanodiscs can be used to systematically study how phosphorylation barcodes regulate arrestin. Those barcodes likely produce different functional outcomes depending on the conformational states of the receptor TM core, which will be important to address in the future. The planar nanodisc platform captures defined lipid composition but does not address membrane curvature, lateral heterogeneity, or higher-order organization, all of which may influence arrestin complex stability^84,85^. Future studies using curved liposomes or more complex membrane systems will be important to test how membrane topology modulates arrestin activation.

Together, our findings support the notion that cooperative control of arrestin activation is governed by a phosphorylation barcode presented by the receptor and a lipid barcode displayed by the membrane. This model prevents accidental activation of arrestins by soluble proteins bearing similar phosphorylation barcodes. It also allows arrestins to operate across compartments with distinct lipid identities and explains why sustained signaling from endosomes may require both specific phosphorylation patterns and compatible membrane environments^86,87^.

## Acknowledgement

We thank Dr. V. V. Gurevich (Vanderbilt University) for providing the Arr2 WT plasmid and Dr. A. Inoue (Kyoto University) for providing NanoBiT constructs. We also thank Dr. Kathryn Schultz for her technical assistance with DEER data collection. This study was supported by the Center for Electron Microscopy (iCEM) at Indiana University School of Medicine. We thank T. Klose and F. Vago at the Purdue Cryo-EM Facility for their technical assistance.

## Author contributions

Q.C. and Y.A. conceptualized this study. Y.A. and Q.C. purified the Arr2 variants and performed the FRET and pull-down assays. Y.Z. performed the DEER experiments and Y.Z., C.K. and A.M. analyzed the DEER data. Y.A. performed the cell-based NanoBit assay and the genome-wide phosphopeptide pattern analysis. Q.C prepared the cryo-EM sample and collected the data. Q.C., Y-Z. Y. and C-L. C. processed the cryo-EM data and performed the modeling. Q.C. and Y.A. wrote the manuscript and all other authors reviewed and edited the manuscript.

## Data and material availability

All data needed to evaluate the conclusions in the paper are available in the main text or the supplementary materials. Additional data related to this paper are available upon reasonable request from the authors. The map and model of the nanodisc–Arr2–ACKR3_pp2_–Fab7 complex have been deposited into the Electron Microscopy Data Bank and Protein Data Bank under accession codes EMD-76522 and PDB ID 12LK, respectively.

## Extended Data

**Extended Data Figure 1.**
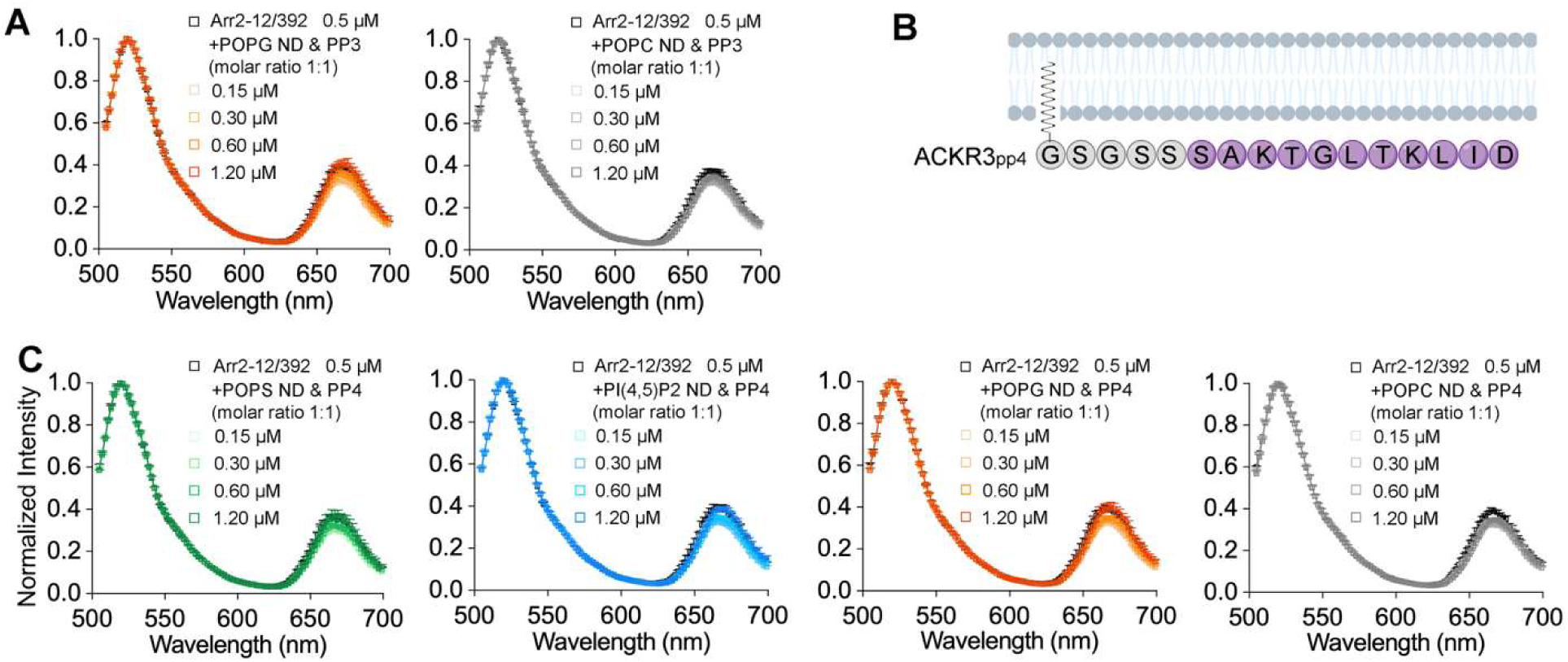
Unphosphorylated nanodisc-tethered peptide does not induce Arr2 activation. **A.** Representative FRET emission spectra of Arr2-12/392 in the presence of a lipidated but non-phosphorylated ACKR3-derived peptide, ACKR3_p4_ (panel B), at increasing concentrations. Spectra are normalized to the donor peak. **B.** Schematic of N-terminally palmitoylated but non-phosphorylated ACKR3_p4_ control peptide tethered to the membrane. **C.** FRET emission spectra of Arr2-12/392 in the presence of ACKR3_p4_ and nanodiscs of different lipid compositions. Peptide and nanodisc concentrations at a 1:1 ratio were titrated over the indicated concentration range.

**Extended Data Figure 2.**
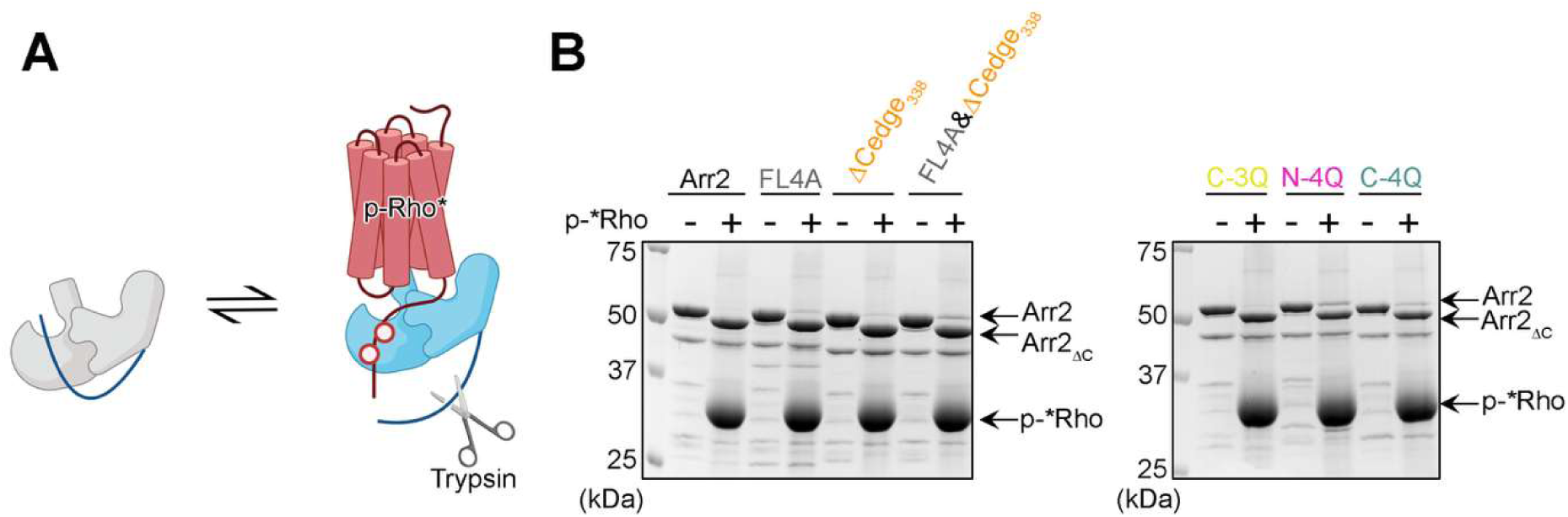
Arr2 variants get activated upon binding to p-Rho*. **A.** P-Rho* binding triggers the release of the autoinhibitory C-terminus of Arr2, rendering it accessible to trypsin digestion and resulting in a truncated form of Arr2 (Arr2_ΔC_). **B.** All the Arr2 variants tested in the Fig. 3E show p-Rho*-dependent release of the C-terminus, suggesting that they are folded correctly and retain functionality.

**Extended Data Figure 3.**
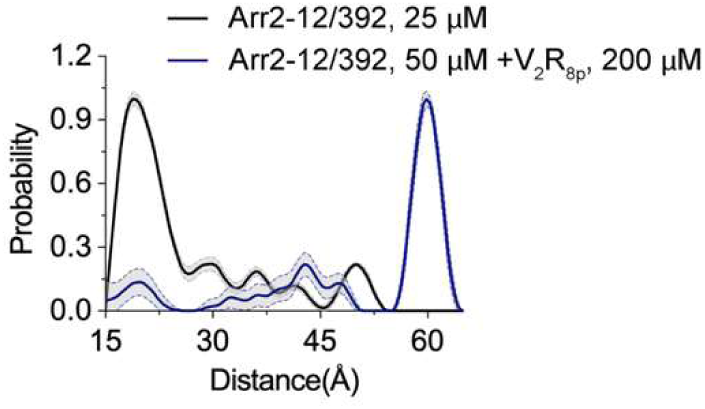
V_2_R_8p_ can induce full Arr2 activation in the absence of membranes. DEER distance distributions of Arr2-12/392 in the absence or the presence of V_2_R_8p_. The distance shifts from ∼19Å in the basal state to >55 Å in the presence of V_2_R_8p_, consistent with C-tail release, indicating Arr2 activation.

**Extended Data Figure 4.**
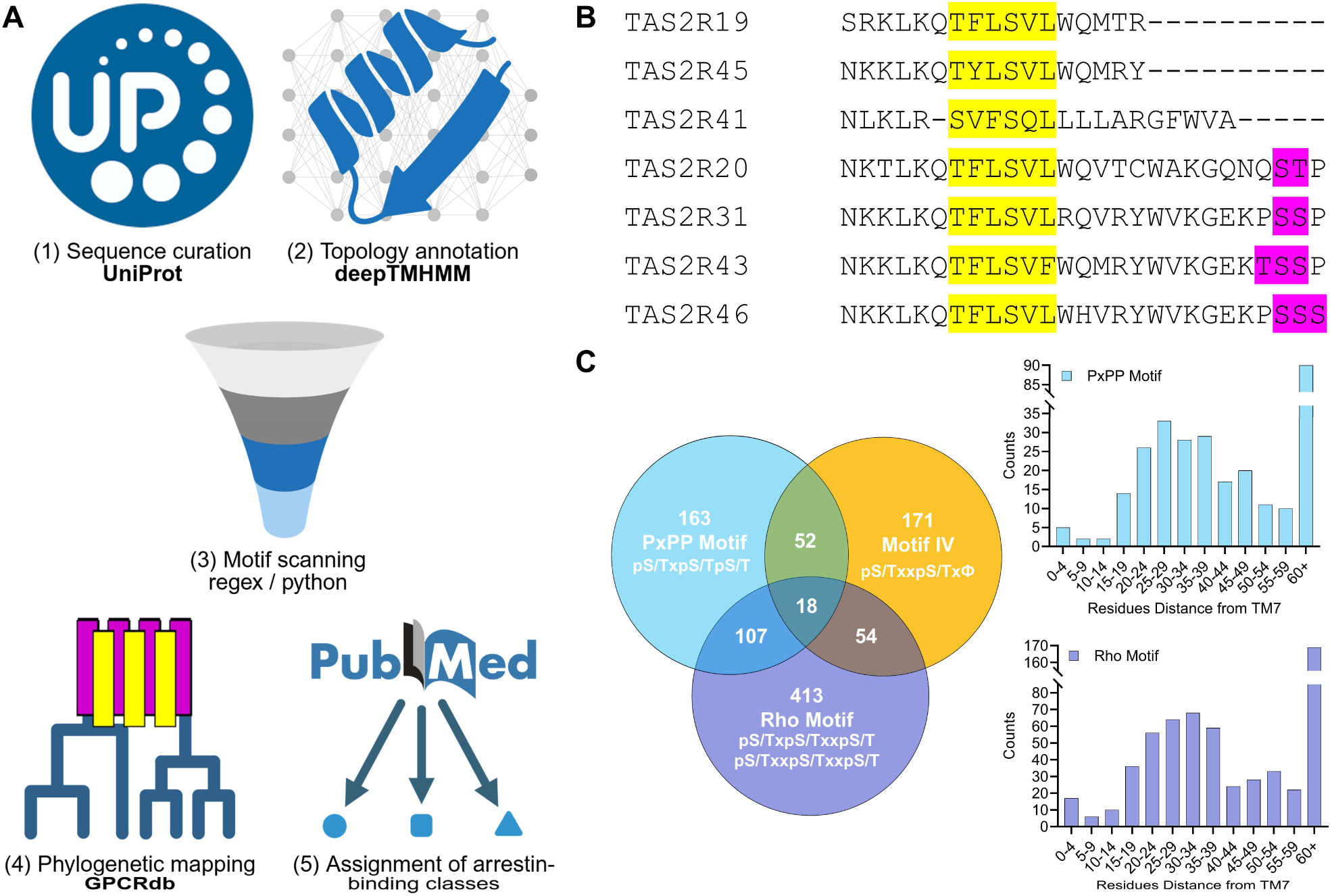
Computational workflow and comparative analysis of arrestin-binding phosphorylation motifs across the GPCRome. **A.** Schematic of the computational pipeline used for global motif analysis. Human GPCR sequences were curated from UniProt, annotated for transmembrane topology using DeepTMHMM, scanned for phosphorylation motifs using regex-based pattern matching, mapped onto GPCR phylogeny using GPCRdb, and cross-referenced with published literature to assign arrestin-binding classifications. **B.** Sequence alignment of C-terminal sequences of select Tas2R family members, including high-affinity Motif II (yellow) and additional distal phosphorylation sites (magenta). **C.** Overlap and positional distribution of Motif IV with previously described arrestin-binding motifs, pS/TxpS/TpS/T (“PxPP”) and pS/Tx(x)pS/TxxpS/T (“Rho”). Venn diagram shows shared and unique occurrences of each motif across GPCR sequences. Histograms depict the distance of motif occurrences from TM7, binned by starting position, highlighting the enrichment of arrestin-binding motifs in membrane-proximal regions.

**Extended Data Figure 5.**
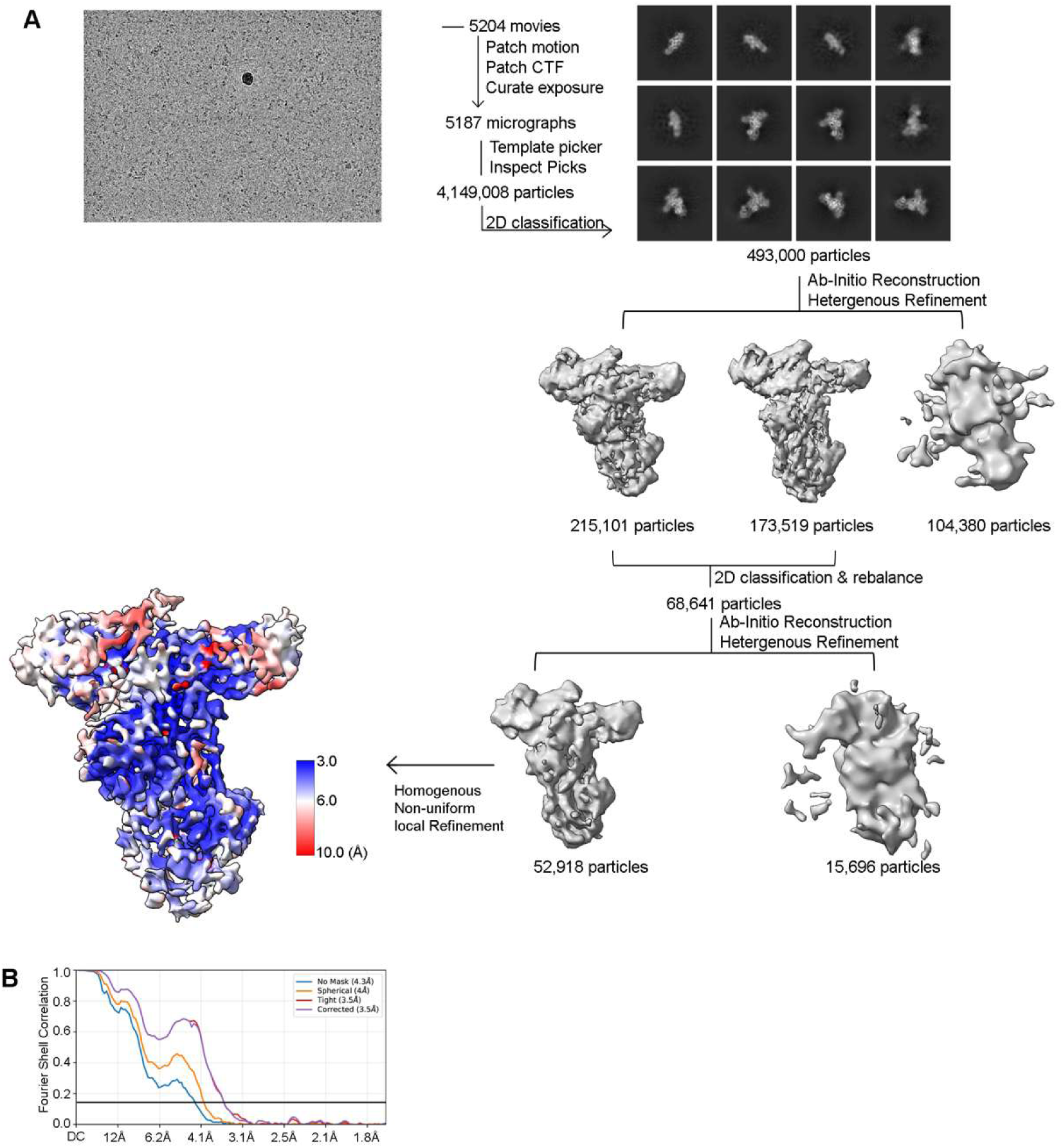
Workflow of cryo-EM data processing and resolution analysis of the nanodisc–Arr2–ACKR3_pp2_–Fab7 complex. **A.** Representative micrograph shows well-distributed complexes. The cryo-EM data processing workflow is shown. Local resolution estimation is calculated by cryoSPARC. **B.** FSC curves calculated by cryoSPARC with 0.143 as a cutoff.

## Methods

### Expression and Purification of Arrestins

Expression and purification of Arr2 WT and variants from *E. coli* were performed using two different protocols. Arr2 WT, Arr2-FL4A, Arr2-ΔCedge_338_, Arr2-FL4A+ΔCedge_338_, Arr2* and Arr2-12C/392C purifications were performed as described previously with several modifications^88^. Briefly, the pTrcHisB plasmid containing the bovine Arr2 was transformed into *E. coli* Rosetta BL21 cells and protein expression was induced with 25 µM IPTG for 4 hours at 30°C. The pelleted bacteria were resuspended and homogenized in buffer containing 20 mM MOPS (pH 7.5), 200 mM NaCl, 5 mM EDTA, 2 mM DTT, 1 mM PMSF, leupeptin, and lima bean trypsin protease inhibitor. Cells were lysed using a microfluidizer and the lysate was clarified by centrifugation at 18,000 rpm for 30 minutes. Arrestin was precipitated from the supernatant by adding (NH_4_)_2_SO_4_ to a final concentration 0.32 g/ml and the precipitate was collected by centrifugation at 18,000 rpm for 30 minutes. The pelleted precipitate was dissolved in a buffer containing 20 mM MOPS (pH 7.5), 2 mM EDTA, and 1 mM DTT, and centrifuged at 18,000 rpm for 30 minutes to remove the insoluble fraction. The supernatant was loaded onto a 5 ml HiTrap Heparin HP column (Cytiva) and eluted with a linear NaCl gradient (0.3-0.8 M for Arr2 and mutants, 0.4-0.9 M for Arr2* and Arr2-12/392). Fractions containing Arr2 were identified by SDS-PAGE followed by staining with InstantBlue® Coomassie Protein Stain (Abcam) and combined. The salt concentration of the pooled fractions was adjusted to 100 mM and then loaded onto a 5 ml HiTrap Q HP column (Cytiva), a 5 ml HiTrap SP column (Cytiva) and a 5 ml HiTrap Heparin HP column (Cytiva). Most contaminants bound to either the Q or the SP column. Arr2 WT and mutants flowed through the Q and SP columns and bound to Heparin column. After uncoupling the columns, Arr2 was eluted with a linear NaCl gradient (0.3-0.8 M) from the heparin column. Arr2* and Arr2-12/392 bound to the SP column and were eluted with a linear NaCl gradient (0.1-0.5 M) from the SP column. The arrestin-containing fractions identified via SDS-PAGE and Coomassie staining were concentrated with a 30 kDa cutoff Amicon concentrator (Millipore) and further purified using a Superdex 200 increase 10/300 GL column (Cytiva) equilibrated with 20 mM MOPS (pH 7.5), 150 mM NaCl, and 0.5 mM TCEP. For Arr2-12/392C, during the size exclusion chromatography purification step, 0.2 mM DTT instead of 0.5 mM TCEP was used to avoid interference with the following labeling reaction. The combined peak fractions were concentrated with a 30 kDa cutoff Amicon concentrator (Millipore), and stored at −80 °C.

Arr2-N-4Q, Arr2-C-3Q and Arr2-C-4Q failed to express using the protocol described above. We therefore followed the procedures described previously^89^ with some modifications. This construct contains an N-terminal octahistidine tag, followed by a 3C protease site and a GGSGGS linker. The Arr2 mutant sequences were codon-optimized for expression in *E. coli* and cloned into a pET-15b vector. All constructs were synthesized by Genescript. The plasmid was transformed into *E. coli* Rosetta BL21 cells and grown in TB at 37 °C until the OD600 reached 1.0. Temperature was reduced to 25 °C and cells were induced with 25 µM IPTG when the OD600 reached 2.0. Cells were harvested after overnight induction. The pelleted bacteria were resuspended and homogenized in buffer containing 20 mM MOPS (pH 7.5), 500 mM NaCl, 10% glycerol, 5 mM 2-mercaptoethanol (β-ME), 1 mM PMSF, leupeptin, and lima bean trypsin protease inhibitor. Cells were lysed using a microfluidizer and the lysate was clarified by centrifugation at 18,000 rpm for 30 minutes. The supernatant was loaded onto a 5 ml HisPur^™^ Ni-NTA Resin (Thermo) and washed with 100 ml of buffer containing 20 mM MOPS (pH 7.5), 500 mM NaCl, 20 mM imidazole, 5% glycerol, 5 mM β-ME and then 50 ml of buffer containing 20 mM MOPS (pH 7.5), 100 mM NaCl, 40 mM imidazole, 5% glycerol, 5 mM β-ME. The column was then eluted with buffer containing 20 mM MOPS (pH 7.5), 100 mM NaCl, 200 mM imidazole, 5% glycerol, 5 mM β-ME in 5 ml fractions. Fractions containing Arr2 were identified by SDS-PAGE followed by staining with InstantBlue® Coomassie Protein Stain (Abcam) and combined. The concentration was measured using Nanodrop and 3C protease was added at 1:100 (w/w) Arr2 protein to cut the His tag overnight at 4 °C. EDTA was added at a final concentration of 2 mM to limit non-specific protease digestion of Arr2. The mixture was loaded onto a 5 ml HiTrap Heparin HP column (Cytiva) the next day and was eluted with a linear NaCl gradient (0.4-0.9 M). The arrestin-containing fractions identified via SDS-PAGE and Coomassie staining were concentrated with a 30 kDa cutoff Amicon concentrator (Millipore) and further purified using a Superdex 200 increase 10/300 GL column (Cytiva) equilibrated with 20 mM MOPS (pH 7.5), 150 mM NaCl, and 0.5 mM TCEP. The combined peak fractions were concentrated with a 30 kDa cutoff Amicon concentrator (Millipore), and stored at −80 °C.

### Fluorophore labeling of arrestin for FRET measurement

Double Cys variants of Arr2 (Arr2-12C/392C) with all native cysteines mutated were expressed and purified as described above with the following modifications. Right before labeling, purified Arr2-12C/392C protein was loaded onto a 5 ml HiTrap Desalting column (Cytiva) equilibrated with 20 mM MOPS (pH 7.5) and 150 mM NaCl to remove the reducing agent. AlexaFluor 488 C_5_ Maleimide (Thermo) was added at a 1:1 molar ratio to Arr2-12C/392C. ATTO 647N (AAT Bioquest) was added at a 3.5:1 molar ratio to Arr2-12C/392C. The mixture was incubated at 4°C overnight in the dark. The labeled sample was loaded onto a 5 ml HiTrap Desalting column (Cytiva) equilibrated with 20 mM MOPS (pH 7.5) and 150 mM NaCl to remove excess fluorophore. The fractions containing labeled arrestins were collected, concentrated with a 30 kDa cutoff Amicon concentrator, and stored at −80°C.

### Expression and purification of NW9

Expression and purification of NW9 from *E. coli* cells were performed as described previously^90^. Briefly, the pET28a plasmid containing NW9 was transformed into *E. coli* Rosetta BL21 cells and protein expression was induced at an OD_600_ of 0.6 using 1 mM IPTG for 3 hours. The pelleted bacteria were resuspended and homogenized in buffer containing 50 mM HEPES (pH 8.0), 500 mM NaCl, 1% Triton X-100, 1 mM PMSF, leupeptin, and lima bean trypsin protease inhibitor. Cells were lysed using a microfluidizer and the lysate was clarified by centrifugation at 12,000 rpm for 30 minutes. The supernatant was collected and loaded twice at room temperature onto 10 ml HisPur^™^ Ni-NTA Resin (Thermo) equilibrated with 20 mM HEPES (pH 8.0) and 500 mM NaCl. The column was washed sequentially with: (I) 100 ml of 20 mM HEPES (pH 8.0), 500 mM NaCl, and 50 mM sodium cholate; (II) 100 ml of 20 mM HEPES (pH 8.0), 500 mM NaCl, and 50 mM imidazole; and (III) 100 ml of 20 mM HEPES (pH 8.0), 500 mM NaCl, and 100 mM imidazole. NW9 was eluted with 100 ml of 20 mM HEPES (pH 8.0), 100 mM NaCl, and 300 mM imidazole. Fractions containing NW9 were identified by SDS-PAGE and Coomassie blue staining. Combined fractions were concentrated with a 30 kDa cutoff Amicon concentrator and loaded onto a 5 ml HiTrap Desalting column (GE) equilibrated with 50 mM HEPES (pH 8.0) and 100 mM NaCl at 500 µl at a time. Fractions containing NW9 were concentrated to ∼10 mg/ml and stored at − 80°C.

### Preparation of Nanodisc

All lipids were purchased from Avanti Polar Lipids. A lipid-to-NW9 ratio of 60:1 was used to assemble empty nanodiscs. All lipids were dissolved in chloroform except for the brain PI(4,5)P_2_, which was dissolved in a mixture of chloroform, methanol and water. Lipids were mixed at desired ratios and dried under a stream of N_2_ gas. The vials were then placed in a desiccator for at least 30 minutes to remove residual solvent. Lipids were solubilized using 10% sodium cholate. NW9 and biobeads (Bio-Rad) were added to the solubilized lipids, and the mixture was incubated on a rotator overnight at 4°C.

### Preparation of hyper-phosphorylated rhodopsin

The isolation of rod outer segments (ROS) and the preparation of hyper-phosphorylated rhodopsin were performed as described previously^91^. In brief, the bovine retina was homogenized and resuspended in 25.5% sucrose. It was then overlaid on two sucrose layers containing 32.25% and 27.125% sucrose. The sucrose gradient was centrifuged in a swinging bucket rotor (SW-27) at 18,000 rpm for 90 minutes. ROS settled between the two layers and were resuspended in buffer containing 20 mM HEPES (pH 8.0) and 2 mM MgCl_2_. All procedures were conducted in a dark room under red light.

For phosphorylation, 2 μM of ROS was incubated with 100 nM GRK1 in a buffer containing 50 mM HEPES (pH 8.0), 500 μM ATP, and 10 mM MgCl_2_ under light at room temperature for one hour. After phosphorylation, the membrane was washed twice with 20 mM HEPES (pH 8.0) and 100 mM NaCl and then incubated with 10-fold molar excess of 11-cis retinal in the dark at room temperature for one hour for regeneration.

### Trypsin digestion

Arr2 WT and mutants at 5 µM were incubated with 15 µM p-Rho* in buffer containing 20 mM MOPS (pH 7.5), 100 mM NaCl, and 20 mM CaCl_2_ at room temperature for 10 minutes. Trypsin at a final concentration of 4 μg/ml was added to initiate digestion at room temperature for 5 minutes. The reactions were quenched with 4x sample loading buffer, run on SDS-PAGE, and stained with InstantBlue® Coomassie Protein Stain (Abcam).

### FRET measurement

The labeled Arr2 at 0.5 μM was incubated with various concentrations of the nanodiscs and phosphopeptide mixture at a 1:1 molar ratio. The reactions were incubated at room temperature for 10 minutes and then transferred to a 384-well black plate (Corning) for FRET measurement using a plate reader (BioTek Synergy H1). The excitation wavelength was set to 476 nm and the emission was monitored from 485 nm to 700 nm at a step size of 2 nm.

### DEER Spectroscopy

The double-Cys Arr2-12/392 variant was incubated with a 40-fold molar excess of 4-maleimido-TEMPO (MAL-6, Sigma–Aldrich) overnight at 4 °C under gentle agitation. Excess spin labels were removed by dialysis with buffer containing 50 mM MOPS (pH 7.0) and 100 mM NaCl.

The labeled Arr2 at 25 µM was incubated with 80 µM p-Rho* (**Fig. 1D**), or 250 µM ACKR3_pp3_ with or without 100 µM nanodiscs (**Fig. 1D, F**). For V_2_R_8p_, the labeled Arr2 at 50 µM was incubated with 200 µM peptide (**Extended Data Fig. 2**). DEER data were collected at 50 K using a Q-band Bruker ELEXSYS 580 and an overcoupled EN5107D2 resonator as previously described^7^. Briefly, samples containing 20% deuterated glycerol as a cryoprotectant were held in quartz capillaries and flash-frozen in a dry ice and acetone mixture. Raw data were phased, background-corrected, and analyzed using the LongDistances1077 software program written by C. Altenbach (University of California, Los Angeles, CA).

### Nanodisc pull-down assays

For arrestin pull-down assays, each binding assay reaction was set with a total volume of 25 µl containing 4 µM nanodisc, and 5 µM Arr2 variant. For the reactions including the peptides, each binding assay reaction was set with an additional 8 µM peptides. The nanodisc and arrestin were incubated in the binding buffer containing 20 mM MOPS (pH 7.5) and 100 mM NaCl for 20 minutes at room temperature. For each reaction, 3 µl of Ni-NTA magnetic beads (PureCube) were added and incubated on a rotator for 60 minutes at room temperature.

The beads were washed three times with 1 ml of the binding buffer and eluted with 20 µl of the binding buffer supplemented with 200 mM imidazole. The eluted proteins were run on SDS-PAGE and stained with InstantBlue® Coomassie Protein Stain (Abcam). The densities of arrestins and NW9 were quantified using Image Lab software (BioRad), and the ratio of bound arrestins to NW9 on the same gel was calculated to represent relative binding.

### Peptide synthesis

All the ACKR3 peptides and V_2_R_pp1_ and V_2_R_pp2_ were synthesized by the Peptide 2.0 Inc. The V_2_R_8p_ was synthesized by Tufts University and is a generous gift from Ziarek lab at Northwest University.

### Cell culture and DNA transfection

HEK293 cells (ATCC) were grown in 100 mm dishes (Fisher Scientific) in DMEM (high glucose, GlutaMax), supplemented with 10% (v/v) fetal bovine serum, 100 U/ml penicillin, 100 mg/ml streptomycin. Cells were cultured in a humidified incubator under 5% CO_2_ at 37°C. On the day before transfection, cells were plated in 6-well dishes (Fisher Scientific) at a density of 5×10^5^ cells per dish. At 60-80% confluency, the cells were transfected using PEI (Fisher Scientific).

Arr2 recruitment to GLP1R and PTH1R was measured using NanoBiT complementation assay as described previously^30,92^, with some modifications. HEK293T cells were seeded at 6×10^5^ in 6-well plates in 2 ml complete DMEM. After 24h, a mixture of 0.3 µg GPCR-SmBiT, 0.3 µg LgBiT-Arrestin, and 2.4 µg empty pcDNA3.1 plasmids was transfected using 3 µl PEI/1 µg DNA. After 24h, media was aspirated and the cells were washed with PBS and detached using 0.1 ml of Trypsin/EDTA for 2 min followed by resuspension in complete DMEM. The cells were seeded at ca. 4×10^5^ using 100 µL of cell suspension per well into a white 96-well plate (Alkali Scientific). After 18-24h, cells were washed with 90 µl warm assay buffer (HBSS + 5 mM HEPES) and 90 µL of warm substrate solution was added (5 µM coelenterazine in assay buffer). The baseline luminescence was recorded with a microplate reader equilibrated to 37°C (Molecular Devices iD5). 10 µl of a 10x ligand serial dilution in assay buffer + 0.1% (w/v) BSA were added and luminescence was measured for 15 min. Luminescence values of stimulated wells at saturation (8-10 min) were averaged, divided by the baseline, and normalized to vehicle-treated wells.

### Global sequence analysis

FASTA sequences of all human GPCRs were downloaded from the UniProt database ^93^ and subjected to batch-wise structure prediction by DeepTMHMM ^94^ to assign intracellular loop segments. This sequence database was subjected to regex-based search for putative phospho-motifs, Motif I-IV described here, and pS/TxpS/TpS/T, and pS/TxpS/TxxpS/T or pS/TxxpS/TxxpS/T (Rho motifs). Distinct occurrences of matches were counted and unique receptors containing one or more motifs were reported and visualized as a phylogenetic tree using the Data Mapper feature of GPCRdb ^56,95,96^. The distances of the first residue of each motif to the end of TM7 as predicted by DeepTMHMM were binned and visualized using Prism. All related preprocessing and analysis code is available under https://github.com/Yasmin361/phosphomotifs.

To assign arrestin coupling classes, manual literature search was conducted for each receptor using PubMed. Experimental evidence from fluorescence imaging or BRET-based recruitment, trafficking and colocalization assays was evaluated. Receptors were classified as Class B if sustained arrestin association or co-trafficking with arrestins to endocytic compartments was reported, Class A if arrestin interactions were transient or rapidly reversible, and uncoupled if no arrestin recruitment was detected under the reported experimental conditions. When explicitly stated by the authors, published classifications were adopted. Receptors lacking sufficient experimental evidence were assigned as unknown, and in cases of conflicting data, most recent studies were weighted more heavily.

### Cryo-EM sample preparation

5 µM Arr2 was incubated with 5 µM 10% PI(4,5)P_2_ nanodiscs, 5 µM Fab7 and 50 µM ACKR3_pp2_ at room temperature for 20 minutes before freezing. Quantifoil R1.2/1.3 300-mesh Cu grids were glow-discharged using EasiGlow at 25 mA for 60 seconds. The complex (3.3 µl) was applied to the grids and the grids were blotted with filter paper for 6 seconds with blot force 2 and 100% humidity before being plunge-frozen in liquid ethane using a Vitrobot MK IV (Thermo Fisher Scientific). Data were collected on a Titan Krios G4 electron microscope (Thermo Fisher Scientific) equipped with a post-GIF K3 direct electron detector (Gatan) in the Purdue Life Sciences Cryo-EM Facility. Micrographs were collected in super-resolution mode with a pixel size of 0.411 Å, at a defocus range of 0.6 to 2.5 µm using EPU, and 40 frames were recorded for each movie stack at a frame rate of 78 milliseconds per frame and a total dose of 59.5 electrons/Å^2^.

### Cryo-EM data processing

Cryo-EM movies were imported to cryoSPARC v4.7.1 and processed using the standard workflow^97^. Beam-induced motion was corrected and binned two-fold using Patch Motion in cryoSPARC. The contrast transfer function (CTF) parameters were estimated using the Patch CTF module. Blob picker was used to pick particles on a small set of micrographs to generate class averages as templates for subsequent autopicking using template picker. Several rounds of 2D classification were performed to exclude bad particles that fell into 2D averages with poor features. Particles from different views were selected to generate three initial models using *ab initio* reconstruction which were refined using the heterogeneous refinement. The two classes with reasonable 3D reconstructions were grouped and subjected to another round of 2D classification. The 40 well-defined 2D classes, containing 372,173 particles, were regrouped into 7 classes, and particle numbers were rebalanced using “cryoSPARC Rebalance 2D Classes” with a rebalance factor of 0.6, yielding 68,614 particles for subsequent ab initio model building, heterogeneous refinement, homogeneous refinement, non-uniform refinement and local refinement^98^. The processing flowchart is shown in **Extended Data Fig 5**.

### Model building and refinement

The cryo-EM structure of active Arr2 in complex with GRK5-phosphorylated ACKR3 and Fab7 (PDB entry 9E82) was modified to omit the receptor core density and the lower half of the Fab7, and then docked into the cryo-EM map using Phenix v1.2.2.2-5419^99^. The resulting model was further improved through real space refinement in Phenix v1.2.2.2-5419 and manual adjustment in COOT v0.9.8.96^100^. Figures were prepared using PyMOL v3.3.1^101^ and ChimeraX v1.8^102^.

